# Yeast elongation factor homolog New1 protects a subset of mRNAs from degradation by no-go decay

**DOI:** 10.1101/2024.11.26.625534

**Authors:** Max Müller, Lena Sophie Tittel, Elisabeth Petfalski, Kaushik Viswanathan Iyer, Stefan Pastore, Tamer Butto, Marie-Luise Winz

## Abstract

New1 is a homologue of the essential yeast translation elongation factor eEF3. Lack of New1 has previously been shown to induce queueing of ribosomes upstream of the stop codon on mRNAs encoding specific C-terminal amino acids, primarily lysine and arginine. Here, we used UV crosslinking and analysis of cDNA, long-read nanopore sequencing and proteomics to address the open question of what consequences such queues have for the yeast cell. We show that these queues are ribosomal collisions, which are recognized by the collision sensor and E3 ubiquitin ligase Hel2, marking these collided ribosome complexes for mRNA degradation via canonical no-go decay. Decay is initiated by Cue2-mediated cleavage upstream of the stop codon. Ultimately, this leads to downregulation of encoded proteins, including highly abundant and important metabolic enzymes Pgk1 and Gpm1, as well as translation elongation factors eEF1-alpha and eEF1-beta. Collisions and resulting downstream effects are codon-, rather than amino acid dependent. E.g., for C-terminal lysine and arginine, only specific codons induce collisions upon lack of New1. Our study shows that New1 protects highly abundant and essential genes from degradation by no-go decay that would otherwise occur in the absence of translation inhibitors or other direct perturbations of translation.

## INTRODUCTION

Translation of genetic information from mRNA into proteins by ribosomes is a highly conserved key process in all living organisms. While translation normally proceeds from the start codon ‘AUG’ to a stop codon (typically ‘UAA’, ‘UAG’, and ‘UGA’), many different problems can occur during translation elongation. Those potentially lead to stalling of ribosomes, and ensuing collision of subsequent ribosomes into the leading, stalled ribosome. This way collided di-ribosomes (disomes) or higher-order polysomes are formed (1–3). Reasons for ribosome stalling and hence, collisions can be found at the levels of mRNAs, tRNAs and ribosomes themselves and due to translation inhibitors, such as cycloheximide, anisomycin and other antibiotics. At the mRNA level, translation roadblocks can occur as stable secondary structures (4), damage, such as oxidation and alkylation (5, 6), truncation (7), as well as poorly translatable codon sequences. For example, poly(A), coding for poly(lysine), is difficult to translate in both, yeast and mammals due to the specific structure during accommodation of the mRNA in the ribosomal A-site, which impedes binding of the aminoacylated lysine tRNA (8, 9). On the other hand, in yeast, one of the arginine codons (CGA) is decoded slowly due to poor base pairing and low abundance of its decoding tRNA-R(ICG) (9, 10). Codon-based stalling is thus governed by both, features of mRNA and tRNA. In addition, lack of specific (aminoacylated) tRNAs, e.g., during amino acid starvation, can also lead to stalling (11). Finally, features of the ribosome, such as faulty rRNA, as well as translation inhibitors can also cause stalling and collisions, triggering quality control (3, 12–14). Ribosome stalling not only causes the production of potentially deleterious truncated proteins, but ribosome collisions can also induce frameshifting (15, 16). Therefore, different translation quality control pathways have evolved, as reviewed in (17, 18). While ribosome-associated quality control (RQC) degrades truncated nascent peptides (19), no-go decay (NGD) removes stalling-prone mRNAs (4). The E3 ubiquitin ligase Hel2 (ZNF598 in mammals) is central to collision sensing and triggering of RQC and NGD (2, 16, 20, 21). This protein binds to the disome interface formed by the small subunits of collided ribosomes. This interface contains two entities of ribosome-associated protein Asc1 (RACK1 in mammals) (2, 3). Hel2 polyubiquitinates ribosomal proteins uS10 (or eS10 in mammals), as well as uS3. This ubiquitination mark helps recruit the RQC trigger (RQT) complex, consisting of Slh1, and ubiquitin binding proteins Cue3 and Rqt4 in yeast (20, 22, 23) (ASCC3, ASCC2 and TRIP-4/ASC-1(20, 24, 25), forming the ASC-1 complex with ASCC1 in mammals). The helicase Slh1 splits leading ribosomes in collisions in an ATP-dependent manner (20, 23, 25–27). The mechanism has been proposed to rely on Slh1 pulling on the mRNA that protrudes from the mRNA entry channel of the leading ribosome, whereby the colliding ribosome acts as a wedge to split the small from the large subunit (23). Both, Cue3 and Rqt4 bind to K63-linked polyubiquitin (28), thereby facilitating recruitment to collided ribosomes, and bind to and stabilize the Slh1 structure (23), but cause only mild defects in splitting ribosomes when lacking individually (20, 24, 25, 29). However, lack of both at the same time causes defects resembling those seen upon *SLH1* deletion (20). Since Slh1 requires an mRNA overhang for its activity, the RQT complex cannot split ribosomes that are stalled at the 3’-end of the transcript, where the entry channel is not occupied by mRNA. Splitting of such ribosomes requires Dom34 (PELOTA in mammals) and Hbs1, which structurally resemble the canonical release factors of translation termination, eRF1 and eRF3 (30–32). For splitting, a third factor, Rli1 (ABCE1 in mammals) is required.

As previously reviewed (18, 17, 33–42), splitting yields a free small subunit, as well as a peptidyl-tRNA stuck in a large subunit. This peptide will be degraded by RQC. At the same time, the mRNA in a collision can also be targeted for degradation, by NGD. Here, several pathways have been described, where the canonical NGD pathway relies on ribosomal protein polyubiquitination by Hel2 (2, 21), and endonucleolytic cleavage by Cue2 (26). An RQC-coupled, and an RQC-uncoupled pathway have been described (2, 43). RQC-coupled NGD relies on ribosome splitting by Slh1, followed by endonucleolytic cleavage, which was first reported to take place in the A-site of the collided ribosome (2). However, cleavage between A- and P-site, as well as in the E-site have been apparent in a more recent report (43). On the other hand, splitting is dispensable for RQC-uncoupled NGD, where cleavage occurs upstream of the collided ribosome, and ribosomal protein eS7 is first monoubiquitinated by Not4, and then polyubiquitinated by Hel2 (2, 43). Upon endonucleolytic cleavage, the resulting 3’-fragment is degraded 5’-to-3’ by exonuclease Xrn1, whereas the 3’-fragment is degraded by the RNA exosome, with help from the Ski complex (4). While the original publication (4) described the splitting factor Dom34 to be essential for canonical NGD, Cue2-dependent cleavage was shown to still occur in the absence of Dom34 (26, 44). Although canonical, endonuclease-based NGD was described first, it appears that the majority of mRNAs bearing collided ribosomes are degraded 5’-to-3’ by Xrn1, without the need for Cue2 (26), but potentially with the help of two additional factors, Smy2 and Syh1 (45).

We were interested in identifying novel players in translation quality control. Inspecting genetic interactions with the genes encoding Hel2 and other NGD factors in yeast *S. cerevisiae*, we identified the *NEW1* gene. *NEW1* displayed positive genetic interactions (characterized by decreases in fitness that are lower in double deletants than expected, judging by the effect in single deletants) with *HEL2, CUE2, DOM34, XRN1* and *SKI3* (encoding part of the Ski complex, needed for exosome function). Indeed, for *HEL2*, the positive genetic interaction was highest with *NEW1*, compared to all other genes in the dataset (46). On the other hand, *NEW1* displayed negative genetic interactions (fitness loss greater than expected) with all three genes encoding yeast RQT factors (Figure 1). New1 is an ABCF protein (47) with close homology to fungal specific and essential translation elongation factor eEF3, encoded by the *YEF3* gene, but which contains an additional N--terminal prion domain (48, 49). eEF3 has been shown to be involved in ribosome translocation, where it facilitates dissociation of the E-site tRNA (50) and supposedly also in A-site tRNA selection (51, 52). Furthermore, eEF3 stimulates translation termination by eRF1/eRF3 and its lack was reported to cause stop codon readthrough (53). Finally, it was also shown to participate in the recycling of post-termination complexes (54). In comparison, less is known about the roles of the eEF3 homolog New1. New1 was shown to associate with polysomes (55). Lack of New1 had been shown to cause a cold sensitivity phenotype, as well as a ribosome biogenesis defect (56), but has also been reported to cause ribosomal queuing on C-terminal arginine (R) or lysine (K) codons, suggesting a role in translation termination and/or ribosome recycling (55). Interestingly, overexpression of New1 can rescue the growth defect caused by eEF3 depletion to some extent, but conversely, overexpression of eEF3 cannot rescue defects caused by lack of New1 (55), suggesting that this protein can fulfil several functions, one of which is specific to New1.

**Figure 1:**
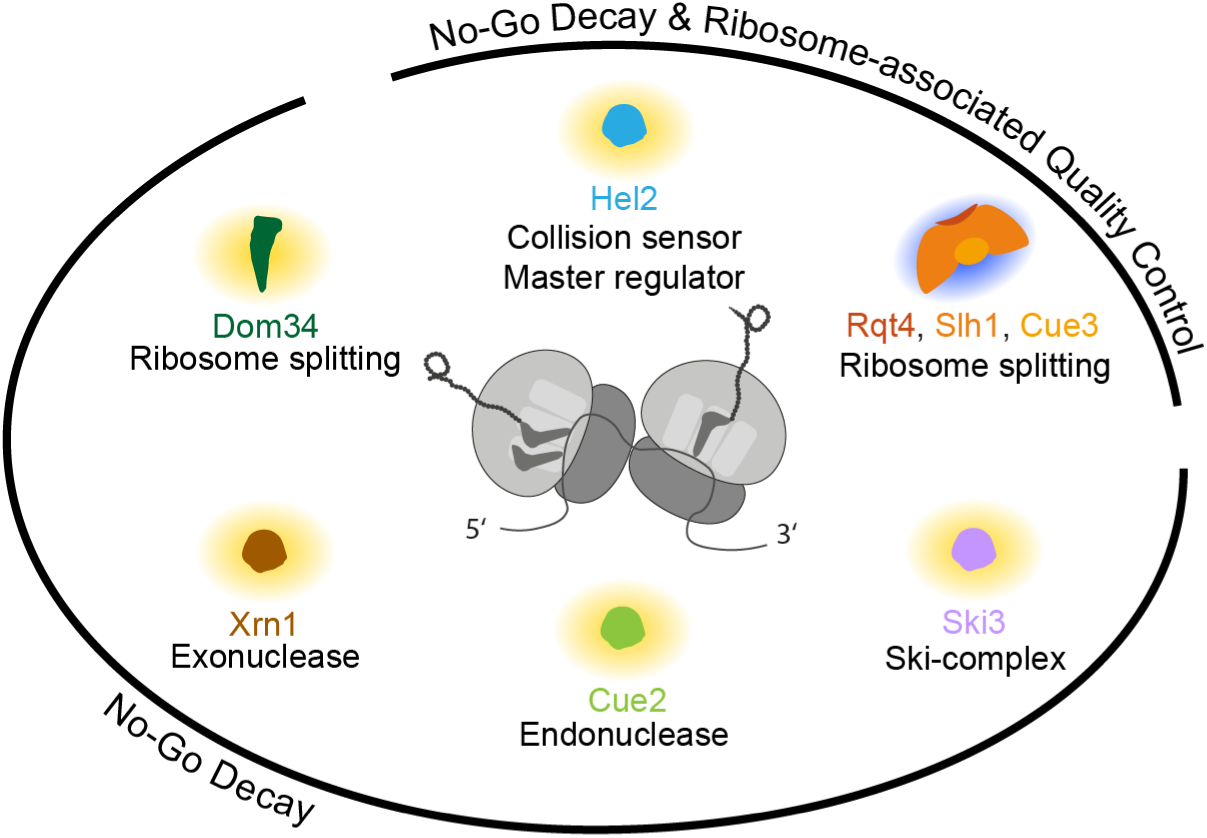
No-go decay (and ribosome-associated quality control) factors encoded by genes that exhibit positive (yellow halo) or negative (blue halo) genetic interactions with *NEW1*, and their roles in no-go decay.

Taking into account the genetic interactions with translation quality control factors, in this work we use a combination of biochemical experiments, UV crosslinking and analysis of cDNA (CRAC) (57), (long-read) RNA sequencing, and proteomics to study effects of *NEW1* deletion on translation quality control. We discover that ribosomal queues in the absence of New1 represent ribosome collisions that occur at specific C-terminal codons. Further, we show that these collisions are sensed by the translation quality control trigger factor Hel2, triggering canonical NGD, of affected mRNAs, initiated by Cue2-mediated cleavage upstream of the stop codon. Several of these mRNAs encode important metabolic enzymes like phosphoglycerate kinase 1 (Pgk1). Consequently, we demonstrate that the gene products of affected mRNAs also tend to be reduced at the protein level. Finally, we proceed to show that New1 associates not only with ribosomal RNA, but also with mRNA, with a preference for 3’-ends, and with tRNA. Our results indicate that New1 protects specific mRNAs from ribosomal collisions and ensuing NGD, with crucial effects on yeast proteostasis.

## MATERIAL AND METHODS

### Strains, oligonucleotides, plasmids and other materials

All yeast strains, oligonucleotides, plasmids, other materials and devices utilized in this project are listed in Supplementary Tables S1-S5, respectively.

### Construction of Plasmids

#### New1 overexpression plasmid

For the construction of a New1 overexpressing plasmid, the coding sequence encoding New1-FLAG was amplified from genomic DNA of a genomically, C-terminally tagged New1-FLAG-TEV-ProteinA2 expressing strain, with Phusion high fidelity polymerase and oligonucleotides harboring XbaI restriction sites as 5’-overhangs. The PCR product was cleaned up via size selection, followed by overnight XbaI digestion at 37°C (0.0075 U/µL XbaI, 1*x* Tango, 50-75 ng/µL DNA). The plasmid pKK148 (58) was digested in the same way. To prevent backbone self-ligation, 0.05 U/µL Fast AP was added to an aliquot of the digested backbone and incubated for 25 min at 37°C. After heat inactivation at 75°C for 10 min, size selection was performed for both PCR product and plasmid backbone via agarose gel extraction. The ligation was conducted with a backbone to insert ratio of 5:1 for the New1-FLAG overexpression vector and non-dephosphorylated backbone was used for empty vector ligation for 1 h at room temperature (1x Ligase buffer, 0.075 U/µL T4 DNA Ligase). The ligation product was transformed into chemically competent DH5ɑ *E. coli* cells and selected on 100 µg/mL ampicillin. Colonies were restreaked and validated via PCR. Plasmids were extracted and the coding sequence of *NEW1-FLAG* was sequenced by Sanger -sequencing (Eurofins Genomics, Mix2Seq), as well as the ligated ends of the empty vector.

#### FLAG-TEV-ProteinA2 tagging plasmid

For His6-TEV-ProteinA2 tagging pBS1539-HTP (57) was used as template for site-directed mutagenesis cloning. pBS1539-HTP was amplified with high fidelity polymerase excluding His6 sequence with 5’-phosphorylated oligonucleotides, bearing the FLAG sequence. Ligation, transformation, and validation were performed as described above.

#### Readthrough-reporter

The coding sequence of GPF was amplified from pGTRR (21, 29) with a forward primer including a XbaI restriction site. The reverse primer was used to manipulate the identity of the C-terminal GFP codon, either to keep the native lysine (AAA) codon or to mutate it to an AAG codon. Furthermore, a stop codon (UAA) was introduce followed by a glycine spacer (GGA) and 0-2 adenosine nucleotides to encode in all reading frames downstream the stop codon the 3x-FLAG-tag. The PCR product was size selected via agarose gel extraction and used for a second amplification with the same forward primer and a second reverse primer to complete the 3x-FLAG sequence and insert an aspartic acid spacer (GAT) to avoid a proceeding lysine (AAG) codon in front of the 3x-FLAG stop codon (UAA). This reverse primer also includes another XbaI digestion site. The remaining cloning was performed as described for the construction of the New1 overexpression vector. As a control construct, the C-terminal GFP codon was omitted, and the 3x-FLAG-tag was placed in the same reading frame as the GFP sequence.

### Construction of strains

Strains were cultured in YPD, at 30°C and 220 rpm. Gene deletion or tagging, as well as plasmid transformation were performed using the “LiAc/SS Carrier DNA/PEG Method” (59) Genes were replaced by either KanMX (using pFA6a-kanMX (60) or pYM18 (61)) or HphMX (using pyM20 (61) cassette or the LEU2 gene (YEp181-CUP1-His-Ubi (62)) and validated via PCR.

Tagging of genes was performed by amplifying the tagging cassette from pBS1539-HTP or pBS1539-FTP with Phusion high fidelity polymerase and oligonucleotides containing 5’-overhangs which anneal to the 5’- or 3’-UTR of the corresponding gene of interest, respectively. Tagging was validated via PCR and Western Blot (see Western Blot section, grown at 30°C), probed with peroxidase anti-peroxidase soluble complex antibody (1:2, 000) over night at 4°C in 5% w/v milk PBST.

### Crosslinking and analysis of cDNA

Crosslinking and analysis of cDNA (CRAC) was performed as previously described (21)] with the following differences: For 20°C culturing, -URA-TRP media was used, whereas for 30°C culturing, -TRP media was used. All samples were crosslinked in a Vari-X-Link (UVO3), with a UV 254 nm dose of 120 mJ (0.7 L (30°C), or 1.0 L (20°C)). For 30°C samples, all further steps were followed as previously described (21). For 20°C samples the following differences in library preparation applied: For purification of tagged proteins magnetic beads were used. Magnetic IgG beads were prepared from 2 mL tosyl activated M280 Dynalbeads, by removing supernatant on the magnet, washing three times with 2 mL 0.1 M sodium phosphate, pH 7.4, mixing by vortexing and removing supernatant. 200 µl rabbit IgG (14 mg/mL in milliQ water) and 800 µl 0.1 M sodium phosphate, pH 7.4 were mixed with the beads by vortexing, then 700 µl of 3 M ammoniumsulphate were added, mixed by vortexing, and incubated overnight at 37°C. Upon removal of supernatant, 2 mL PBS, 0.5% Tween, pH 7.4 and 200 µl BSA (10 mg/ml in milliQ water) were mixed with the beads and incubated for 1 h at 37°C. Supernatant was removed and 2 ml PBS, 0.1% Tween, pH 7.4 were added and washed (rotating) three times for 5 min. Beads were resuspended in 1.2 mL PBS, 0.1% Tween. 100 µl of beads were used for Hel2-HTP samples, 150 µl of beads were used for New1-HTP and untagged samples. Magnetic IgG beads were washed twice prior to immobilization with 2 bead volumes of lysis buffer. RNase digestion was performed after on-bead TEV cleavage (for elution from IgG beads) in 730 µl volume of TEV cleavage solution with 1:100 diluted RNaceIT for 5 min (1.3 µl RNaceIT: 30°C samples), 4.5 min (1 µl RNaceIT: 20°C, New1-HTP samples and untagged controls) or 4 min (1 µl RNaceIT: 20°C, Hel2-HTP samples). For 20°C samples, additional changes were applied: Following SDS gel electrophoresis on NuPAGE Bis-Tris 4-12% gels, the transfer step was skipped. Instead, gel pieces were excised, and protease digestion was performed with crushed gel pieces, pooling several samples in one tube. Reverse transcription was performed with Superscript III, with indexed RT-primer (30°C samples) or non-indexed RT primer (20°C samples). For 20°C samples, RNA:cDNA hybrids after reverse transcription were additionally subjected to RNaseH cleavage for 30 min at 37°C, whereas RTs with indexed RT-primer (30°C samples) were treated with Exonuclease I. PCRs (performed with LATaq and PCR primers as mentioned in Supplementary Table S3) were either purified with Qiagen PCR purification kit (30°C samples) or (20°C samples) remaining primers were removed from PCR products by digestion with Exonuclease I (1 to 1.5 µl in 150 µl PCR mixture), and DNA was phenol extracted and ethanol precipitated, instead of kit-based purification. 20°C sample PCR products were size-selected on TBE polyacrylamide gels and eluted in milliQ water and ethanol precipitated, whereas 30°C samples were size selected on Metaphor agarose (Lonza), as previously described (21). All samples (30°C, 2 biological replicates, 1 sequencing run per replicate; 20°C, 2 biological replicates, 1 sequencing run per replicate) were sequenced on a MiniSeq system (Illumina), using 75 bp single-end sequencing. For 30°C samples, index reads were also sequenced. Barcodes are given in Supplementary Table S6.

### Western Blot

Cultures were grown at 20°C and 220 rpm shaking in SDM drop out medium corresponding to the harboring plasmid marker or in SDM complete media. Secondary cultures were inoculated at OD600 0.05 (10 mL) and grown until log phase, reaching an OD600 of 0.5-0.9, harvested by centrifugation including an additional washing step with ultrapure water and pellets were stored at -20°C. Pellets were lysed by a chemical lysis approach (63), including an additional centrifugation step (14, 000 rpm, 5 min) before loading 10 µL of the lysate. 10% polyacrylamide gels were run for 15 min at 75 V, followed by 100 V until the ladder fully resolved. Proteins were transferred onto nitrocellulose membrane for 75 min at 100 V. The quality of the samples and the transfer was checked using Ponceau stain and blocked with 5% w/v milk. Membranes on which Pgk1 was to be quantified were probed overnight at 4°C with anti-alpha Tubulin antibody (1:10, 000) and 2 h at room temperature with anti-Rabbit IgG HRP antibody (1:5, 000) in 5% w/v milk PBST. After washing, membranes were visualized with sufficient ECL solution on the FUSION Pulse TS (Vilber) system. Membranes were then stripped three times for 15 min with mild stripping buffer and washed with PBST. Then, membranes were probed with anti-PGK1 antibody (1:10, 000) antibody at 4°C overnight, followed by 2 h incubation with anti-Mouse (GAM)-HRP conjugate antibody (1:20, 000) in 5% w/v milk in PBST. Signals were quantified using ImageJ and Pgk1 levels were normalized to Tubulin levels. For visualization of the readthrough reporter, anti-FLAG antibody (1:1, 000) or anti-GFP antibody (1:1, 000) was used, both followed by secondary anti-Mouse (GAM)-HRP conjugate antibody (1:20, 000). Pgk1 was probed as described above.

### Northern Blot

Cultures were grown (25 mL) and harvested as described in “Western Blot” section, with the exception that pellets were resuspended in 750 µL TRIzol™ Reagent, snap frozen in liquid nitrogen and stored at -80°C. For RNA preparation, 150 µL zirconia beads were added per sample, vortexed 30 s at 3, 000 rpm and kept on ice for at least 30 s between rounds, in a total of 10 rounds. 150 µL ROTI®C/I was added to each sample, briefly vortexed and kept on ice for 2 min. After centrifugation at 4°C for 10 min at 14, 000 rpm, the aqueous phase was recovered, followed by a second acid phenol/chloroform extraction. RNA was precipitated by adding 1/10 volume 3 M sodium acetate pH 5.2 and 1 volume 2-propanol and samples were kept at -20°C for 1 h. Next, RNA was pelleted at 14, 000 rpm at 4°C for 20 min, washed twice with 70% v/v ethanol, air-dried for 5 min, dissolved in ultrapure water and RNA concentration was measured with the NanoDrop™ 2000 system. Equal amounts of NorthernMax™-Gly Sample Loading Dye and 0.15 µg/µL additional ethidium bromide were added to the samples and incubated 30 min at 50°C. Up to 10 µg total RNA and 2 µL Riboruler low range ladder were loaded on a 1.4% w/v agarose MOPS gel in the Owl™ A5 system. The gel was run with MOPS buffer pH 7 at 50 V for 18-19 h. RNA was transferred using the capillary-based transfer method with 10x SSC pH 7 onto a nylon membrane. Afterwards, RNA was immobilized via crosslinking twice at 1, 200 µJoules × 100 doses with UV Stratalinker™ 1800. The membrane was blocked with hybridization buffer and 2 mg salmon sperm DNA for 1 h at 42-57°C (Hybrid Mini 38, H. Saur). Oligonucleotides were phosphorylated with γ-32P-ATP (1.6 pmol oligonucleotide, 1x reaction buffer A, 1.6 pmol γ-32P-ATP, 0.5 U/µL T4 PNK) for 30 min at 37°C. EDTA pH 8.0 was added to stop the reaction and heat inactivation was performed at 75°C for 10 min. Blots were probed with 5’-32P-phosohorylated oligonucleotides and fresh hybridization buffer overnight. The membrane was washed three times (up to 45 min) with washing solution, and a phosphor screen was exposed to the membrane and visualized with the Typhoon FLA9500 system. If required, membranes were stripped twice for 1 h with 0.5% w/v SDS at 65°C, blocked and re-probed as described before. Signals were quantified with ImageJ.

### Long-read sequencing

Strains were grown, harvested and RNA was prepared as described before (see “Northern Blot” section) with the exception, that cultures were grown in complete supplement mixture (CSM) complete media. After measuring RNA concentration, RNA was stored at -80°C. For strains lacking *SKI2*, 2’, 3’-cyclic phosphate ends (which are the product of NGD mediated endonucleolytic cleavage (7) end healing was conducted with T4 polynucleotide kinase. For this, total RNA was used in a reaction volume of 60 µL total (1x reaction buffer B, 12 µg RNA, 30 U T4 PNK) and incubated for 30 min at 37°C. Afterwards, acid phenol (pH 5.2)/chloroform extraction was performed, followed by precipitation as described before (see “Northern blot” section). Next, poly(A) tailing was conducted according to the manual “Poly(A) Tailing of RNA using *E. coli* Poly(A) Polymerase” (NEB #M0276). After 30 min of 37°C incubation, EDTA was added to stop the reaction and RNA clean-up was performed with RNAClean XP beads at a concentration of 2x. Library preparation and sequencing was performed as described previously (64, 65). An overview of libraries is given in Supplementary Table S7. Samples were sequenced in (biological) triplicate per condition.

### Mass spectrometry

#### Growth and lysis

Cultures were grown (20 mL) as described before (see “Long-read sequencing” section), without adding TRIzol® Reagent. 150 µL RIPA buffer and 150 µL zirconia beads were added to each sample, vortexed at 3, 000 rpm for 30 s, kept on ice for at least 30 s per round, for a total of ten rounds. Supernatant was transferred to a fresh tube and centrifuged at 14, 000 rpm at 4°C for 10 min. Supernatant was transferred to a fresh tube. Lysates were incubated in ice cold sonicator water bath to shear genomic DNA. Protein concentration was measured with ROTI®Quant universal according to manufacturer’s protocol with the plate reader FlexStation (Molecular Devices).

#### Enzymatic protein digestion

All samples were processed using the SP3 approach (66). The proteins were then digested using trypsin overnight at 37°C. The resultant peptide solution was purified by solid phase extraction in C18 StageTips (67).

#### Liquid chromatography tandem mass spectrometry

Peptides were separated via an in-house packed 45-cm analytical column (inner diameter: 75 μm; ReproSil-Pur 120 C18-AQ 1.9-μm silica particles, Dr. Maisch GmbH) on a Vanquish Neo UHPLC system (Thermo Fisher Scientific). The online reversed-phase chromatography separation was conducted through a 100-min non-linear gradient of 1.6-32% acetonitrile in 0.1% formic acid at a nanoflow rate of 300 nl/min. The eluted peptides were sprayed directly by electrospray ionization into an Orbitrap Astral mass spectrometer (Thermo Fisher Scientific). Mass spectrometry was conducted in data-dependent acquisition mode using a top50 method with one full scan in the Orbitrap analyzer (scan range: 325 to 1, 300 m/z; resolution: 120, 000, target value: 3×106, maximum injection time: 20 ms) followed by 50 fragment scans in the Astral analyzer via higher energy collision dissociation (HCD; normalised collision energy: 26%, scan range: 150 to 2, 000 m/z, target value: 1 × 104, maximum injection time: 5 ms, isolation window: 1.4 m/z). Precursor ions of unassigned, +1 or higher than +6 charge state were rejected. Additionally, precursor ions already isolated for fragmentation were dynamically excluded for 20 s.

### Computational analysis

#### Mass spectrometry data processing

Raw data files were processed by MaxQuant software (version 2.1.3.0) (68) using its built-in Andromeda search engine (69). MS/MS spectra were searched against a target-decoy database containing the forward and reverse protein sequences of UniProt *Saccharomyces cerevisiae* reference proteome (release 2023_05; 6, 091 entries) and a default list of common contaminants. Trypsin/P specificity was assigned. Carbamidomethylation of cysteine was set as fixed modification. Methionine oxidation and protein N-terminal acetylation were chosen as variable modifications. A maximum of 2 missed cleavages were allowed. The “second peptides” options were switched on. “Match between runs” was activated. The minimum peptide length was set to 7 amino acids. False discovery rate (FDR) was set to 1% at both peptide and protein levels. For label-free protein quantification, the MaxLFQ algorithm (70) was employed using its default normalization option. Minimum LFQ ratio count was set to one. Both the unique and razor peptides were used for quantification. Differential expression analysis was performed in R statistical environment. Reverse hits, potential contaminants and “only identified by site” protein groups were first filtered out. Proteins were further filtered to retain only those detected in all three replicates in either the wild-type or knock-out group. Following imputation of the missing LFQ intensity values, a linear model was fitted using the limma package (71) to assess the difference between the wild-type and knock-out groups for each protein, with adjustment for multiple testing using the Benjamini-Hochberg approach (72).

#### Sequencing analysis

The stop codon and preceding codons of all open reading frames (ORF) of coding sequences (CDS) (excluding ORFs classified as “dubious” or “pseudogene”) were extracted from “orf_coding.fasta”, downloaded from SGD (http://sgd-archive.yeastgenome.org/sequence/S288C_reference/orf_dna/; accessed 22.07.2024).

#### CRAC data analysis

CRAC data were analyzed, primarily employing the pyCRAC package (73) as previously described (21) for analysis of total hit distributions, tRNA hit distributions, and plotting of mRNA (with deeptools (74), done as previously described for metagene plots that were scaled over part of the gene body) and rRNA read and deletion coverage, always without scaling to library size. For 3’-untranslated region (3’-UTR) binding analysis, deeptools (74) module computeMatrix was used on the same .bigwig files used for analysis of mRNA coverage, but employing a reference .gtf file containing 3’-UTR coordinates instead of ORF coordinates, and using 100 nt upstream and downstream unscaled option, 10 nt bins, and ‘gene-body’ size 50 (so, 5 bins, representing the annotated 3’-UTR). Resulting values for the 5 bins of each individual 3’-UTR were summed up for each sample, and a pseudo-count of 1 was added, to avoid division by zero for cases where no reads were obtained for a 3’-UTR in one of the samples. Then, the ratio of 3’-UTR-sums was calculated for each pair of Hel2-HTP, *new1Δ* and Hel2-HTP, wt sample (replicates 1 and 2 at 30°C, and replicates 1 and 2 at 20°C). For pairs where no 3’-UTR reads were recovered in either sample, the value was set to N/A. Ratios were then normalized to the median of all ratios calculated for a pair of samples, to avoid the influence of factors that influenced mRNA read values (e.g., higher or lower recovery of rRNA vs. mRNA, leading to higher or lower RPM values for mRNA, including 3’-UTR in some samples). Data were then plotted using boxplotR (75) according to either the C-terminal amino acid, or the C-terminal codon.

#### Analysis of long-read sequencing (via Nanopore-Seq)

Data analysis with NanopoReaTA was performed as described previously (65), based on fastq output files from the Oxford Nanopore MinKnow sequencing software. deepTools (74) was used via the command “bamCoverage -bs 1, normalizeUsing CPM –outFileFormat bedgraph” to transform the BAM files into bedgraph files. Based on the gene coordiantes from the gene list, extracted from SGD (and excluding genes harbouring introns), the average sequencing depth for each gene was extracted and the relative fold change of each gene was computed.

#### 3’-end analysis

To extract the 3’-end positions of the aligned sequences from the *ski2Δ* and *new1Δ, ski2Δ* data, the coordinates of each 3’-end were extracted from the corresponding BED file (generated via bedtools) and the number of alignments per positions counted. The counts were extracted for 300 nt upstream of the stop codon coordinate and data from all three replicates were pooled after CPM normalization. The scores for each gene within this 300 nt range were summed up for wt, *ski2*Δ and *new1Δ, ski2*Δ together and the relative value per position was calculated. Published 5Pseq WIG files (76) for wildtype and *new1Δ* were downloaded and 5’-coordinates were extracted for genes of interest, pooled and relative values per position were calculated as described above. Data were smoothed via Gaussian filter approximated to sigma = 1. Additionally, deepTools (74) was used as previously described for CRAC data (21) to generate binned (10 nt bin-size) coverage plots covering 100 nt in front of the start codon, 250 nt behind the stop codon, 250 nt unscaled region after start codon, 400 nt unscaled region in front of the stop codon, and a scaled region of 50 bins in between.

#### Ribo-Seq analysis

Queuing scores (QS) of the already published Ribo-seq data for wt and *new1Δ* strains (55) were computed according to Kasari et al., 2019 (55). Provided Ribo-seq BAM files were also transformed into bedgraph files as described above and analyzed with deeptools (74) similar to CRAC data. For some genes, the QS calculation method did not yield a queuing score. Therefore, in addition, the coverage of the 90 nucleotides (9 bins) preceding the stop codon was divided by the coverage along the entire transcript and log_2_ of the relative fold change between knock-out and wildtype was computed.

#### Generation of box plots

Boxplots were generated using the boxplotR tool (75) with default settings, except for showing data points (jittered), and, in some cases, variable width boxes, logarithmic scaling of y-axis, and adjusting the plot size. 20°C samples are usually shown with blue boxes, 30°C samples with red boxes.

### Western and Northern blot analysis

Western and northern blot signals were quantified with ImageJ (77). Rectangle width was adjusted to average signal size and height of the membrane. Tangents were fitted to the signal specific graphs to determine the area under the curve. Based on the loading control, a normalization factor was computed to adjust the signal of interest and to determine the relative fold change, as well as the log2 fold change.

### Statistical analysis

T-test was performed via Microsoft excel choosing two tailed distributions with an unequal sample variance.

## RESULTS

### The absence of New1 causes recruitment of Hel2 to mRNA 3’-untranslated regions, dependent on the C-terminal codon

Since the absence of New1 had been reported to cause ribosome queuing on mRNAs encoding specific C-terminal amino acids, we wondered whether these queues are equivalent to collided ribosomes and might elicit collision-specific downstream responses, including no-go decay. This does not necessarily need to be the case, as ribosomes can occur as collided disomes and collided higher-order polysomes without eliciting quality control, as exemplified by several disome sequencing studies (78–82).

To find whether this was the case, we tested if the collision sensor Hel2 is recruited preferentially to mRNAs affected by C-terminal ribosomal queuing in the absence of New1. To this end we performed the CLIP-like approach crosslinking and analysis of cDNAs (CRAC)(57), using His_6_-TEV-ProteinA_2_ (HTP) tagged Hel2 in either wt or *new1Δ* strains, at 30°C, and in addition at 20°C, since the lack of New1 causes an increased cold sensitivity (56). We found that, for a subset of mRNAs, Hel2 binding to the 3’-UTR is drastically increased in the absence of New1, at both temperatures (Figure 2A-C, Supplementary Figures S1-S3). Examining closely which parameters determined this specific increase, we found that mRNAs encoding C-terminal lysine (K) or arginine (R) and to a lesser extent cysteine (C), asparagine (N) or serine (S), but not other amino acids, exhibited increased binding of Hel2 to 3’-UTRs (Figure 2A, Supplementary Figure S1), in agreement with observation of queues specifically on such mRNAs (55). Looking more closely, we determined that only specific lysine and arginine codons mediated this effect, the most strongly affected being K(AAA), R(AGG) and R(CGU), but not, e.g., K(AAG) or R(AGA) (Figure 2B). For the mildly affected cysteine, asparagine, and serine, the codon specificity seemed to be less pronounced, but was also more difficult to judge due to lower numbers of observations, especially for cysteine (Supplementary Figure S2). For C-terminal asparagine codons, in particular, the increase in Hel2 recruitment to 3’-UTRs was more pronounced at 20°C than 30°C, in contrast to all other codons (Figure 2A, Supplementary Figure S1). Our findings are in agreement with a recent publication (76), which showed ribosomal queuing for specific C-terminal lysine, arginine, asparagine and serine codons. We therefore conclude that queues indeed represent ribosome collisions that cause recruitment of the collision sensor and E3 ubiquitin ligase Hel2. For all following analyses, codons K(AAA), R(AGG), and R(CGU) will be referred to as ‘strongly affected’ or ‘strong’, all cysteine (C), asparagine (N), and serine (S) codons, as well as arginine codons R(CGA), R(CGC), and R(CGG) will be referred to as ‘mildly affected’ or ‘mild’, and all remaining codons as ‘not affected’ or ‘not’. In the following, most experiments were conducted at 20°C, as the reported phenotypes were stronger at that lower temperature, unless mentioned otherwise.

**Figure 2:**
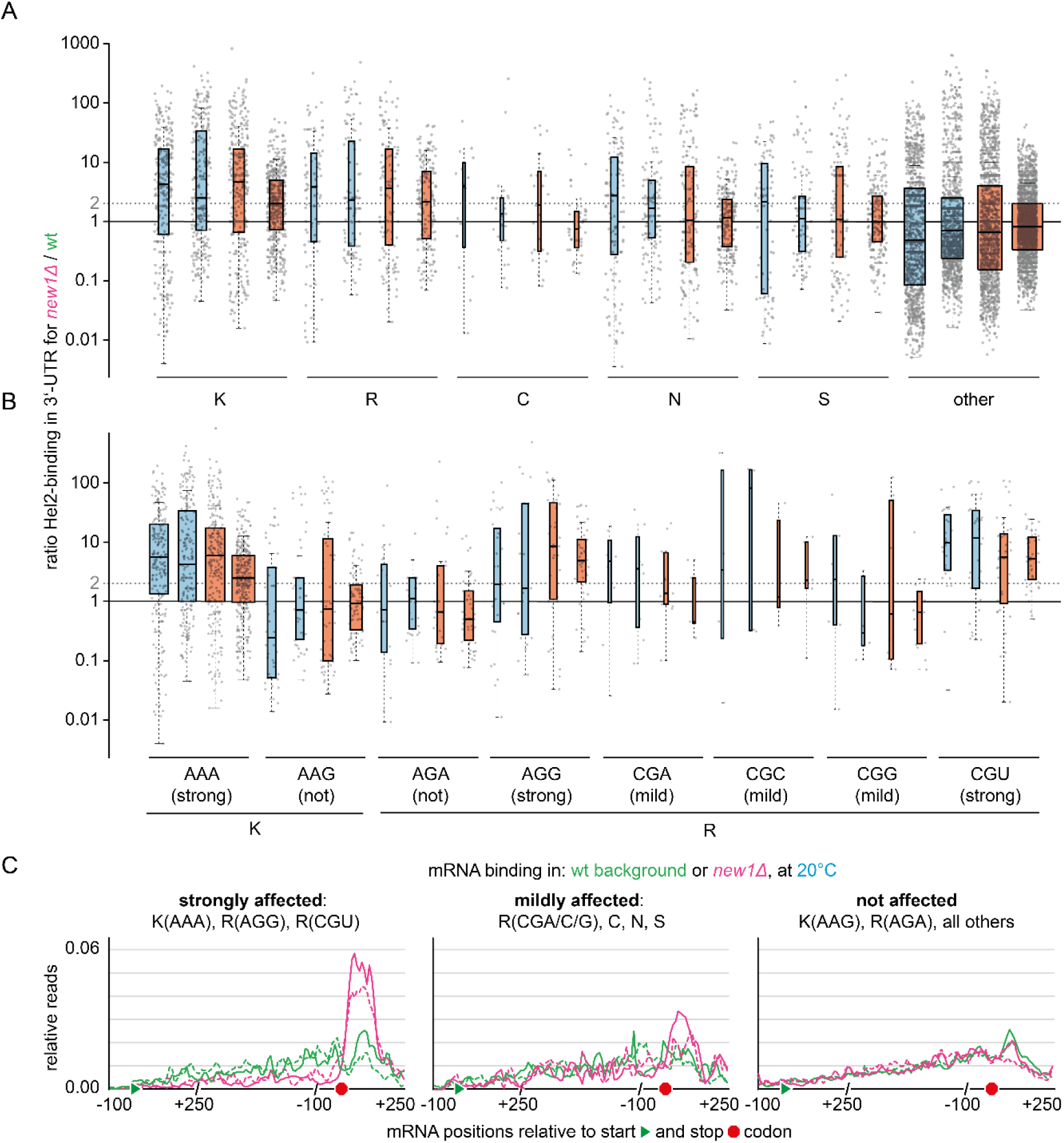
Ratio of Hel2 binding in 3’-UTR for *new1Δ*/wt at 20°C (blue, 2 biological replicates), and 30°C (red, 2 biological replicates) (**A**) for mRNAs encoding C-terminal amino acids lysine (K), arginine (R), cysteine (C), asparagine (N), serine (S) or other (excluding K, R, C, N, S), (**B**) for all lysine and arginine codons. Boxplot description for (**A**) and (**B**): Center lines show the medians, box limits indicate the 25^th^ and 75^th^ percentiles, as determined by R software; whiskers extend 1.5 times the interquartile range from the 25^th^ and 75^th^ percentiles. All data points are represented by dots. Width of the boxes is proportional to the square root of the sample size. (**C**) Metaplots showing Hel2 binding (relative hits) in either wildtype (wt, green, 2 replicates) or *new1Δ* (magenta, 2 replicates) background, at 20°C, for those mRNAs within the group of previously identified top 1000 highest Hel2-bound mRNAs (21) with C-terminal codons being either strongly affected, mildly affected or not affected by lack of New1.

#### Hel2 recruitment to mRNA 3’-UTRs upon lack of New1 is not generally due to stop codon readthrough

Next, we asked why Hel2 binds 3’-UTRs. We argued that this could be due to stop codon readthrough, as reported for lack of eEF3 (53), which could lead to the need of translation quality control, e.g., due to ribosome stalling on poly(A) and other unfavourable sequences (83). We argue that such sequences would be more likely to be present in 3’-UTRs due to lack of evolutionary pressure for codon optimization in untranslated regions. Ribosome densities had been shown to be slightly increased in 3’-UTRs of affected mRNAs, however, stop codon readthrough had previously not been observed using a dual-luciferase readthrough reporter system (55). However, that reporter construct had only been tested in the ‘in-frame’ reading frame. Since ribosome collisions can cause frameshifting (15), we argued that readthrough itself might be caused by frameshifting, as this would lead to part of the stop codon being recognised as a sense codon. In addition, eEF3, the homologue of New1 has been shown to be actively involved in frameshifting at collisions (84). As a consequence, readthrough *c*ould occur in a different reading frame, which was not tested before. Based on this reasoning, we designed a readthrough reporter, employing GFP with a native terminal AAA (strongly affected) or AAG (not affected) codon followed by a ‘UAA’ stop codon, a glycine (GGA) spacer to avoid any potential influence of the stop codon context that might otherwise cause variations (85, 86) between the different reading frame constructs and a 3xFLAG-tag, either in-frame, in the +1 or in the -1 frame (Supplementary Figure S4). The levels of FLAG (and hence readthrough in specific reading frames) were determined by Western blot and compared to a positive control without a stop codon. In our assay, we did not detect significant levels (all samples <0.5% of control, according to FLAG signals) of stop codon readthrough. Even more importantly, we did not observe any differences between wt and *new1Δ*. While writing this manuscript, a report was published (76) which showed that readthrough may occur in the absence of New1, but only under very specific conditions, where the affected C-terminal codon is followed by the weak stop codon ‘UGA’ and an additional nucleotide ‘C’. Other combinations did not allow for marked stop codon readthrough in the dual luciferase assay employed in that study, validating the findings from our reporter assay, in which the affected codon was followed by the strong stop codon ‘UAA’, and followed by the nucleotide ‘G’, generating a strong termination context (85, 86). To exclude that the 3’-UTR enrichment of Hel2 was observed solely due to mRNAs in which the strongly affected codon is followed by the weak ‘UGA’ stop codon and a nucleotide ‘C’, we compared 3’-UTR enrichment in *new1Δ* over wt for this subset (20 mRNAs) to the subset where the strongly affected codon is followed by the stronger stop codons ‘UAA’ or ‘UAG’ and/or a +1 nucleotide ‘not C’ (154 mRNAs for ‘UGA, not C’; 60 mRNAs for ‘not UGA, C’; and 434 mRNAs for ‘not UGA, not C’). 3’-UTR enrichment was observed for all subgroups, although in the ‘UGA, C’ subgroup, it was only detectable at 20°C, but not 30°C (Supplementary Figure S5). Here, it is possible that a certain level of stop codon readthrough on such mRNAs might reduce the number of ribosome collisions, and hence Hel2 recruitment.

Taken together, we conclude that ribosome collisions upstream of stop codons do not generally lead to stop codon readthrough, and that Hel2 mRNA binding in 3’-UTRs is most likely not due to suppression of stop codons. Although this does not fully exclude the possibility that ribosomes might re-initiate translation in the 3’-UTR, as previously observed (78), and that resulting translation problems within the 3’-UTR can lead to Hel2 recruitment, we favour the hypothesis that Hel2 crosslinking in 3’-UTRs is mainly driven by Hel2 binding to collided ribosomes at the stop codon, and making contacts with downstream mRNA. This hypothesis is also supported by the fact that a minor interaction site of Hel2 with the ribosomal small subunit, previously described by us, is localized close to the mRNA exit channel (21), suggesting that Hel2 would also be able to interact with mRNA at and beyond this position.

### Hel2 recruitment upon lack of New1 causes degradation of affected mRNAs by no-go decay

One possible downstream consequence of Hel2-catalyzed ribosomal protein polyubiquitination is NGD. We therefore asked whether mRNAs affected by increased ribosomal queuing upon deletion of *NEW1*, and to which Hel2 is recruited, are targeted by NGD, namely canonical NGD. Canonical NGD involves first endonucleolytic cleavage by Cue2, followed by degradation of endonucleolytic fragments. We decided to probe for both, total mRNA levels and endonucleolytic cleavage fragments. To enable detection of endonucleolytic fragments by Northern blot, we used strains lacking Ski2, a member of the Ski complex, which is a necessary co-factor for the cytosolic RNA exosome (87). As a result, the 5’-fragment resulting from endonucleolytic cleavage is stabilized (4), whereas the 3’-fragment is still degraded by Xrn1. As potentially affected target mRNAs, we chose the highly expressed *PGK1*, *ADH1*, and *GPM1* transcripts, all ending on the K(AAA) codon and exhibiting both, ribosomal queueing and Hel2 3’-UTR recruitment. All three genes encode important metabolic enzymes, involved in glycolysis/gluconeogenesis (*PGK1* and *GPM1*), or alcoholic fermentation (*ADH1*). As a control, we visualized mRNA for another metabolic gene: *TDH3*, which ends on the non-affected alanine (A) amino acid, encoded by the A(GCU) codon. We found that lack of New1 leads to a strong decrease in total mRNA levels (in *SKI2*+ background) of all three affected mRNAs by ∼40-60%, but not the non-affected *TDH3* mRNA (Figure 3A, comparing lanes 1, 2 to lane 3; quantification Figure 3B). This effect is fully rescued by plasmid-based overexpression of C-terminal FLAG tagged New1 (Figure 3A, lane 4; Figure 3B). In the absence of Ski2 and New1, but not Ski2 alone, we observed the appearance of shorter fragments, which we interpret as endonucleolytic 5’-fragments, for all three affected mRNAs, but not the control mRNA of *TDH3*. Endonucleolytic cleavage was suppressed by overexpression of FLAG tagged New1 (Figure 3A, lanes 5-8). For Pgk1, we further tested whether lower mRNA levels translate to decreased protein levels in affected strains. Here, we indeed observed a decrease in Pgk1 protein by approximately 63%, but not in alpha-Tubulin, which ends with the non-affected amino acid phenylalanine (F), encoded by F(UUU): Tub1; or F(UUC): Tub3 (Figure 3C, D). We thus showed for several examples that, in the absence of New1, mRNAs ending with strongly affected codons (here: K(AAA)) are cleaved to form shorter fragments, leading to decreased full-length mRNA levels, supporting the notion that they become targets of NGD. Additionally, our data on Pgk1 suggest that levels of the encoded proteins are also decreased.

**Figure 3.**
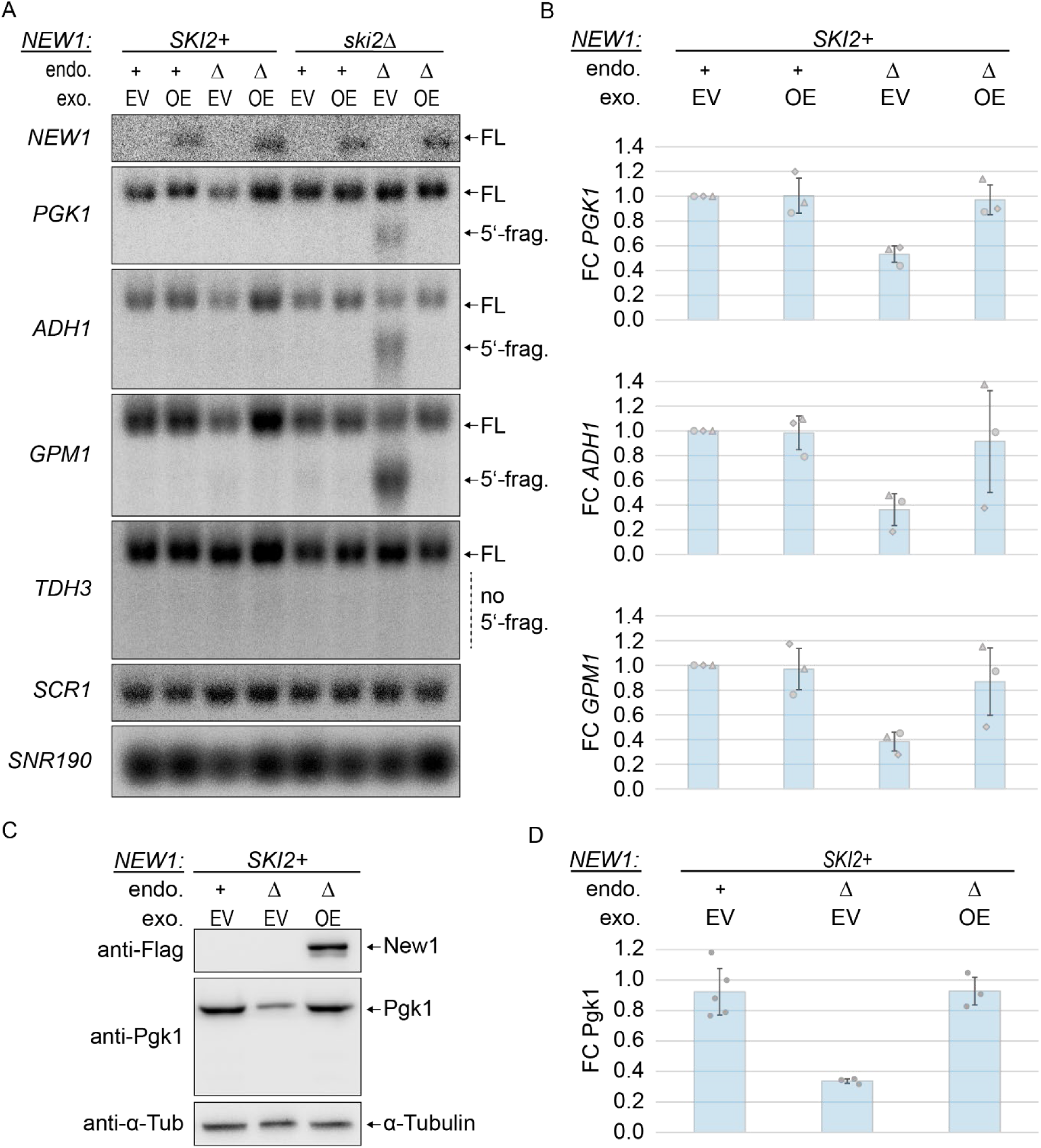
(**A**) Northern blot analysis of full-length (FL) mRNA levels or endonucleolytic 5’-fragment (5’-frag.) for *NEW1*, *PGK1*, *ADH1*, *GPM1*, *TDH3*, *SCR1*, and *SNR190* in *new1Δ* (endo Δ), *NEW1*-positive (endo +) strains, containing an empty vector (exo EV) or New1-overexpression vector (exo OE), and containing or lacking the *SKI2* gene. (**B**) Quantification of full-length mRNA levels for *PGK1*, *ADH1* and *GPM1*, relative to *TDH3* (control), compared to wildtype strains with empty vector (endo +, exo EV), related to (**A**), n = 3 biological replicates. Data were normalized to *SCR1* signals. (**C**) Western blot analysis of Pgk1 levels in *new1Δ* (endo Δ, exo EV), and New1-overexpressing (endo Δ, exo OE) strains, compared to wildtype strains containing an empty vector (endo +, exo EV). (**D**) Quantification of protein levels for Pgk1, relative to Tubulin, related to (**C**), n = 3 biological replicates. Data were normalized to Tubulin signals. Error bars in (**B**) and (**D**) represent 1 standard deviation. Additional replicates shown in Supplementary Figure S6.

#### Degradation of mRNAs upon lack of New1 is caused by canonical no-go decay

Next, we wanted to know whether the putative endonucleolytic cleavage of affected mRNAs depends on known NGD factors and thus represents canonical RQC-coupled NGD. To this end, we generated strains lacking New1 in combination with either of the following known NGD factors: Hel2, Slh1 (helicase of the RQT complex, splits leading ribosomes in collisions, depending on the availability of an mRNA overhang at the ribosomal mRNA entry channel, essential for RQC-coupled NGD (2, 23, 43)), Cue3 and Rqt4 (both ubiquitin-binders, part of the RQT complex (20, 29)), Dom34 (splitting factor for stalled ribosomes without mRNA in the A-site; was reported to be required for NGD (4)) and Cue2 (NGD endonuclease (26)). In the absence of Ski2 (Figure 4A), we only observed *new1Δ*-dependent fragments (Figure 4A) in the presence of Hel2 and Cue2, confirming our interpretation as endonucleolytic cleavage fragments, and consistent with the notion that these factors are essential for canonical NGD (2, 21, 26). For Dom34, we acknowledge conflicting reports on whether or not the factor is required for endonucleolytic cleavage in NGD (4, 26, 44). For *PGK1* mRNA in the absence of New1, however, we were only able to detect endonucleolytic 5’-fragments in its presence.

**Figure 4.**
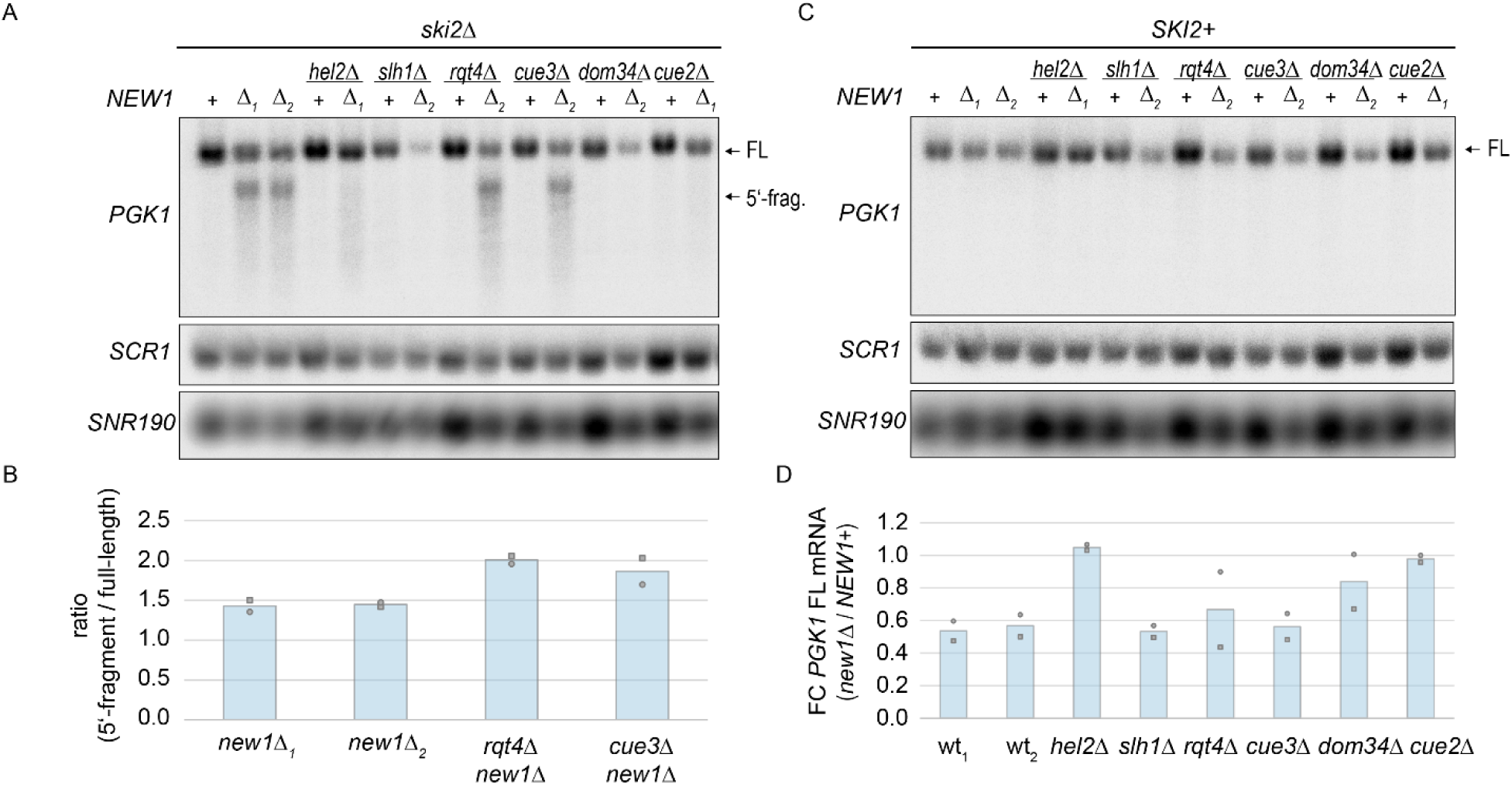
Northern blot analysis of no-go decay dependence on various NGD factors, (**A**) in *ski2Δ* strains, (**B**) quantification for (**A**), (**C**) in *SKI2* containing strains, (**D**) quantification for (**C**). Δ_1_ designates mutants in which the *NEW1* gene was replaced with a KanMX cassette, Δ_2_ designates mutants in which the *NEW1* gene was replaced with a HphMX cassette. In (**D**), wt_1_ designates the sample combination of Δ_1_ and wt, wt_2_ designates the combination of Δ_2_ and wt. Additional replicates shown in Supplementary Figure S7.

We were not able to detect 5’-fragments in the absence of Slh1, the central factor of the RQT complex. This could suggest that NGD for the *PGK1* mRNA in the absence of New1 is RQC-coupled. Surprisingly though, cleavage fragments appeared to be increased in the absence of the two additional factors of the RQT complex, Cue3 or Rqt4 (Figure 4A, B). This may be explained by the fact that Cue3 and Rqt4 are not essential but have a stimulating effect on the splitting function of Slh1 (27). Also, e.g., in case of RQC, the absence of either Cue3 or Rqt4 only causes mild defects, while only the double deletion of both factors causes a phenotype comparable to lack of Slh1 (20, 27, 29). Therefore, in the absence of Cue3 or Rqt4, splitting of leading ribosomes is decreased but still possible. Decreased efficiency of splitting is then likely to enhance queueing, generating more targets for RQC-coupled NGD. In strains containing Ski2, but lacking Hel2, Cue2, or Dom34, we observed a (near) complete stabilization of full-length mRNA after normalization to loading controls (Figure 4C, D). This suggests that, in *new1Δ*, alternative decay mechanisms such as Xrn1-based NGD (26) were not involved in decay of *PGK1* mRNA. In contrast, in the absence of Cue3, Rqt4 and Slh1, we still observed decreased levels of full-length mRNA, comparable to those in the *new1Δ* single knockout strain. Despite the absence of endonucleolytic cleavage fragments in the *new1Δ, ski2Δ, slh1Δ* triple knockout strain, this supports that NGD in *new1Δ* could be RQC-uncoupled. Lack of 5’-fragments in the *new1Δ, ski2Δ, slh1Δ* triple knockout strain could alternatively be explained as a secondary effect of the triple deletion, which also leads to a strong growth defect (data not shown). On the other hand, reduction of full-length mRNA could be due to a reduction of *PGK1* transcription. We also note a smear of shorter-length mRNA observed in the strain lacking New1, Hel2 and Ski2, but not in the strain lacking only New1 and Hel2, suggesting the presence of Hel2-independent decay--. In this strain, however, full-length mRNA was not visibly decreased (Fig. 4A). Future studies may elucidate these phenomena in greater detail. Nonetheless, we can deduce with great certainty that *PGK1* mRNA is targeted to canonical NGD in the absence of New1.

#### Canonical NGD of mRNAs upon lack of New1 is initiated by endonucleolytic cleavages upstream of the stop codon

To obtain high-resolution data on cleavage sites, we performed long-read nanopore sequencing (direct cDNA). In our *SKI2*-deletion strains, the 5’-fragment is stabilized. This fragment would lack the poly(A) tail of the full-length mRNA. Direct cDNA library preparation for nanopore sequencing relies on the use of poly(dT) primers. Thus, we employed *in vitro* polyadenylation to add poly(A) tails also to non-polyadenylated species. However, endonucleolytic cleavage in NGD had been reported to yield 2’, 3’-cyclic phosphates at the 3’-terminus of the 5’-fragment (7), which inhibit polyadenylation. We therefore dephosphorylated RNA with polynucleotide kinase, prior to *in vitro* polyadenylation. Comparing sequencing data for *new1Δ, ski2Δ* double knockout to *ski2Δ* single knockout, we observed the appearance of shorter species of affected mRNAs only for the double knockout (Figure 5A). This was the case for all three previously Northern blot validated *new1Δ*-dependent NGD-targeted mRNAs. Especially for *ADH1*, we observed very heterogeneous fragment lengths, suggesting multiple cleavage sites, between ∼40 and ∼300 nt upstream of the Stop codon (Figure 5B). This size range is similar to previous observations made on stalling reporters (1, 43), and fragment lengths are also in good agreement with previous reports on both RQC-coupled and RQC-uncoupled NGD (43). An approximately 30 nt periodicity (Figure 5B) suggests, that these cleavages result from multiple collisions, also reminiscent of previous reporter data (1). While it is, in principle, possible that these collisions are part of the original queue at the affected C-terminal codons, we are also aware that the 5’-fragment is stabilized in strains that lack Ski2. Hence, translation could be initiated on these fragments, which then represent non-stop mRNAs, leading to further collisions and additional cleavages. This would be similar to what is the case for reporters that contain a self-cleaving ribozyme and, post-cleavage, lack both, stop codon and poly(A) tail, but are still translated (7). Taking into account 5’-phosphate-sequencing data for wt and *new1Δ* strains from a recent publication (76), our data (Figure 5B) suggest that Cue2-cleavage in the first collided ribosome occurs mainly few nt upstream of the E-site, similar to recent findings (43) for RQC-coupled NGD, whereas cleavages within further trailing ribosomes occur close to the 5’-end of ribosome footprints, similar to what is reported for RQC-uncoupled NGD. In contrast, we hardly observed any cleavage sites within the first, stalled ribosome.

**Figure 5.**
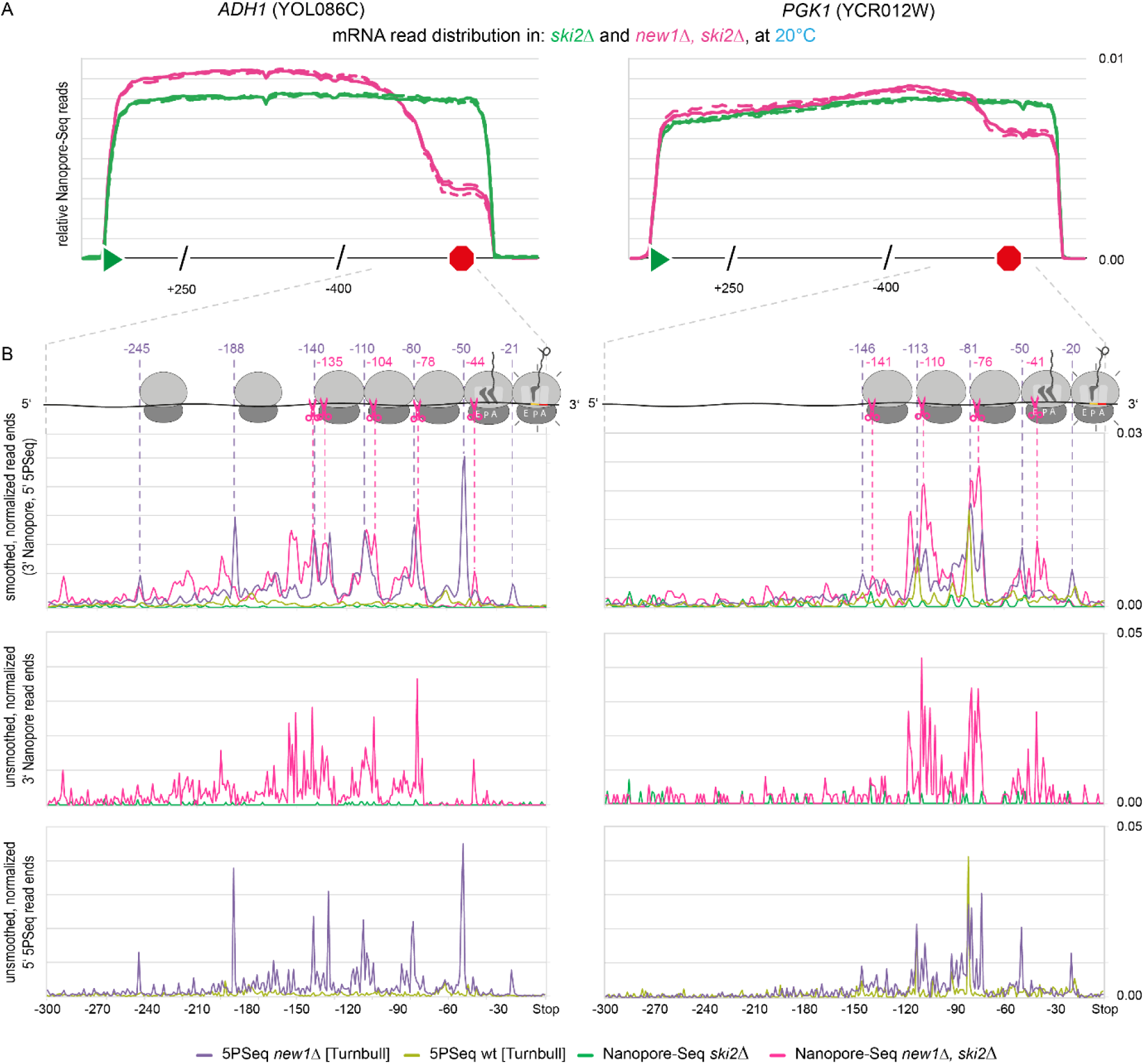
Analysis of endonucleolytic fragments via Nanopore-Seq in *new1Δ, ski2Δ* double knockout and *ski2Δ* single knockout yeast strains. (**A**) Comparison of relative Nanopore-Seq read distribution along the *ADH1* and *PGK1* transcripts. 100 nt upstream of and 250 nt downstream of the start codon, as well as 400 nt upstream of and 250 nt downstream of the stop codon are shown unscaled, whereas the rest of the gene body was scaled. n = 3 biological replicates. (**B**) 3’-end analysis of Nanopore-Seq reads from *new1Δ, ski2Δ* double knockout (magenta, 3 biological replicates pooled) and *ski2Δ* single knockout (green, 3 biological replicates pooled) yeast strains for the last 300 nt of the *ADH1* and *PGK1* transcripts. Data were compared to 5’-ends from 5PSeq reads for *new1Δ* (purple) and wt (light green) representing the position of ribosomes along the mRNA, published by Turnbull et al. (75) Smoothed and unsmoothed data are given. Ribosome positions are schematically shown, based on 5PSeq 5’-ends, and endonucleolytic cleavage sites are indicated by scissors.

In our data, we were able to identify additional targets of endonucleolytic cleavage. These include mRNAs encoding translation elongation factors EF-1 alpha (*TEF1*, *TEF2*; interestingly, EF-1 alpha is a known interactor of New1’s homolog eEF3 (88)), and EF-1 beta (*TEF4*), ribosomal protein S5 (*RPS5*), methyltransferase and ribosomal small subunit processome complex component (*NOP1*), and adenosine kinase (*ADO1*, required for the utilization of S-adenosylmethionine) (Supplementary Figure S8). Except for the Rps5 coding sequence (ending in R(CGU)), all other verified NGD targets contain K(AAA) as the C-terminal codon. For all validated targets, endonucleolytic cleavage sites, as judged by the additional, shorter fragments, were situated about 30-80 nt upstream of stop codons (Figure 5A, Supplementary Figure S8), providing evidence that ribosome collisions that led to NGD did not happen in the 3’-UTR, but rather upstream of stop codons. Although we found that some mRNAs that contain C-terminal K(AAA), R(CGU) or R(AGG) are not targets of endonucleolytic cleavage (e.g., *RPS15*, *RPL41B*) the sequencing depth was not sufficient to analyse most mRNAs that contain affected C-terminal codons. In summary, we show here that NGD upon lack of New1 can trigger canonical NGD for specific and essential genes, which requires Hel2, Cue2 and Dom34.

#### Validated NGD targets are downregulated at the mRNA level

mRNAs that we validated as NGD targets were generally also downregulated at the mRNA level in nanopore sequencing data generated from *new1Δ* strains (containing the *SKI2* gene), compared to wt, employing the standard library preparation protocol without *in vitro* polyadenylation. This was especially true at 20°C, where the only NGD target not down regulated was *NOP1* mRNA (Table 1, Figure 6A). Most of our verified NGD targets were indeed among the most downregulated mRNAs (of mRNAs with CPM>100 to avoid noise, Figure 6A), although the overall effect on all mRNAs containing strongly affected C-terminal codons was not large. This suggests that, as already hinted by our NGD fragment analysis, not all mRNAs affected by C-terminal queueing are targeted to NGD. The finding that *NOP1* levels were not downregulated, despite validated endonucleolytic cleavages in *new1Δ* strains (Supplementary Figure S8) could be due to additional effects that influence RNA levels. One example could be transcriptional upregulation that would counteract downregulation via NGD.

**Figure 6.**
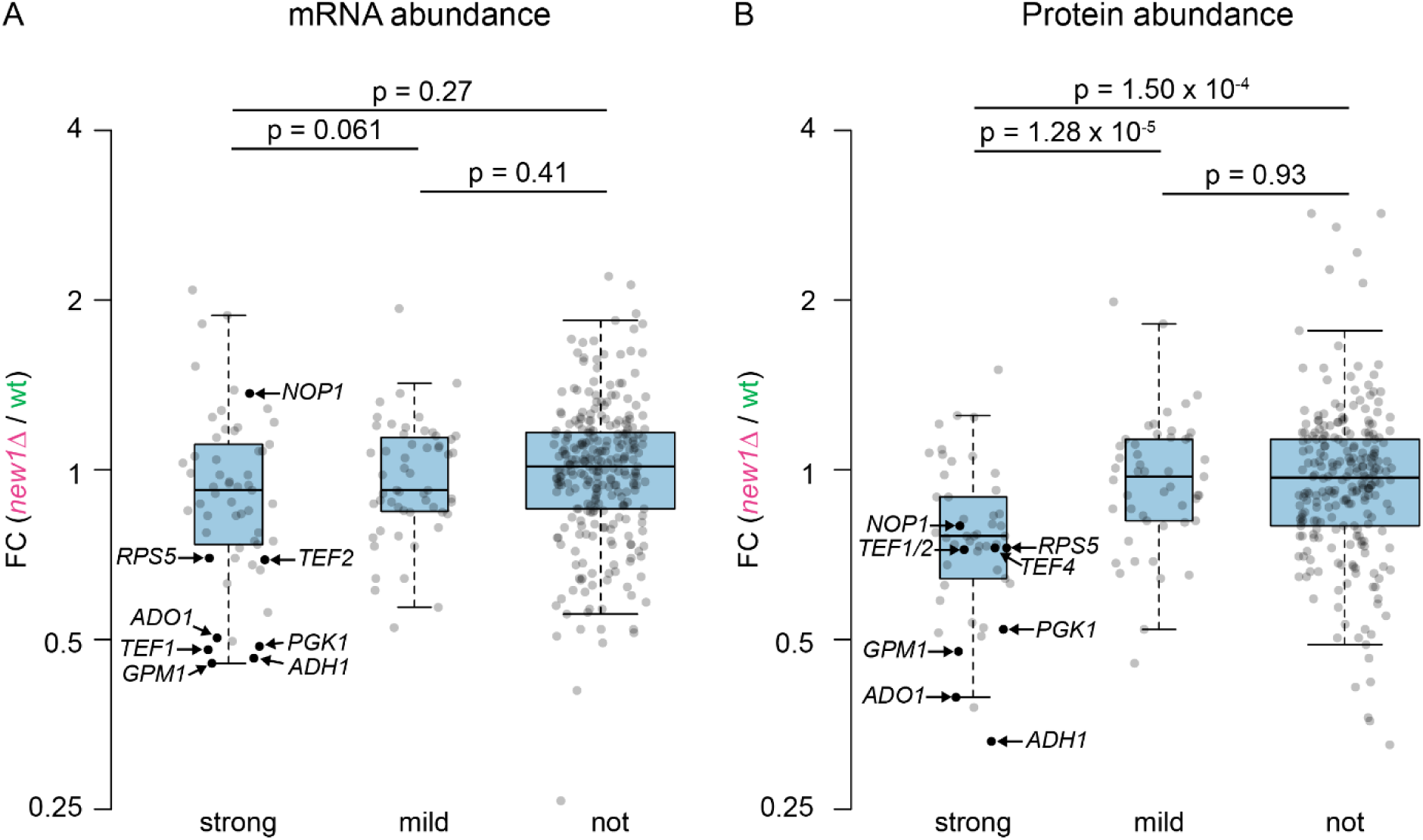
Fold-changes in mRNA, measured by Nanopore-Seq (**A**) and protein, measured by proteomics abundance (**B**) between *new1Δ* and wt at 20°C. Only genes with an overall expression of > 100 CPM (in Nanopore-Seq) are shown to ensure sufficient signal-to-noise ratios. Center lines show the medians, box limits indicate the 25^th^ and 75^th^ percentiles, as determined by R software; whiskers extend 1.5 times the interquartile range from the 25^th^ and 75^th^ percentiles. All data points are represented by dots. Validated NGD targets are highlighted.

**Table 1:** Overview of Hel2 3’-UTR enrichment (log2 fold change (FC) between Hel2 CRAC data from *new1Δ* vs. wt); log2FC of mRNA abundance (*new1Δ* vs. wt); log2FC of protein abundance (*new1Δ* vs. wt), log2FC of ribosome densities upstream of stop codons (based on published data (55), and log2 of queueing scores (QS) calculated as described (55) from two different publications (55, 75). Values are given for all validated NGD targets, as well as three negative controls used in this work: Tubulin (*TUB1* and *TUB3* genes) and Tdh3. *TEF1* and *TEF2* mRNAs encode identical protein products, therefore protein level is only given once. For several genes, QS or ribosome density could not be calculated successfully. Therefore, log_2_ foldchange of ribosome density in the last 90 nt of the coding sequence was calculated based on the same published data from Kasari et al. (54).

| Gene |  |  | log<sub>2</sub>FC(Hel2 3'-UTR CRAC) |  | log<sub>2</sub>FC(mRNA abundance) |  | log<sub>2</sub>FC(protein abundance) | log<sub>2</sub>FC(ribo-some density) |  | log<sub>2</sub>(QS) ref. (55) |  | log<sub>2</sub>(QS) ref. (76) |  |
| --- | --- | --- | --- | --- | --- | --- | --- | --- | --- | --- | --- | --- | --- |
| ID | name | C-term. codon | 20°C | 30°C | 20°C | 30°C | 20°C | 20°C | 30°C | 20°C | 30°C | 20°C | 30°C |
| Validated as NGD targets in absence of New1 |  |  |  |  |  |  |  |  |  |  |  |  |  |
| *YCR012W* | *PGK1* | AAA | 3.36 | 2.38 | -1.04 | 0.19 | -0.94 | 0.9 | 0.74 | 1.97 | 1.52 | N/A | 1.36 |
| *YOL086C* | *ADH1* | AAA | 1.05 | 2.18 | -1.11 | -0.97 | -1.6 | 0.9 | 1.21 | 2.09 | 1.15 | 3.45 | 3.51 |
| *YKL152C* | *GPM1* | AAA | 2.06 | 2.53 | -1.14 | -0.75 | -1.07 | 0.95 | 1.25 | 2.68 | 1.98 | 3.13 | 3.02 |
| *YPR080W* | *TEF1* | AAA | 2.41 | 1.17 | -1.06 | -0.67 | -0.48 | 1.05 | 1.57 | N/A | N/A | N/A | 0.77 |
| *YBR118W* | *TEF2* | AAA | 3.13 | 1.71 | -0.53 | -0.15 |  | 1.04 | 1.54 | N/A | N/A | N/A | 1.02 |
| *YKL081W* | *TEF4* | AAA | 2.64 | 2.47 | -0.2 | -0.39 | -0.46 | 0.92 | 1.19 | N/A | N/A | 3.03 | 2.48 |
| *YJR123W* | *RPS5* | CGU | 2.66 | 0.84 | -0.52 | 0.46 | -0.46 | 0.74 | 1.15 | 1.82 | 1.22 | 1.74 | 1.69 |
| *YDL014W* | *NOP1* | AAA | 7.19 | 1.36 | 0.45 | -0.93 | -0.38 | 0.75 | 1.01 | 2.28 | 1.92 | 0.85 | 1.42 |
| *YJR105W* | *ADO1* | AAA | 2.22 | 3.9 | -0.99 | -0.9 | -1.34 | 0.65 | 0.86 | 1.63 | N/A | 1.92 | 0.78 |
| Negative controls |  |  |  |  |  |  |  |  |  |  |  |  |  |
| *YML085C* | *TUB1* | UUU | -1.38 | -0.37 | -0.14 | 0.69 | -0.17 | -0.04 | 0.1 | N/A | N/A | N/A | -0.64 |
| *YML124C* | *TUB3* | UUC | 2.97 | N/A | 0.01 | 0.43 | -0.39 | N/A | N/A | N/A | N/A | N/A | -0.44 |
| *YGR192C* | *TDH3* | GCU | -1.92 | -1.75 | 0.06 | -0.46 | -0.16 | 0.14 | 0.48 | 0.37 | 0.44 | 0.41 | 0.27 |

### Lack of New1 leads to decreased levels of proteins translated from many affected mRNAs

Now, as we had already observed decreased protein levels for Pgk1 via western blot, matching decreases in *PGK1* mRNA levels, we asked whether NGD triggered in the absence of New1 generally leads to decreased protein levels for affected genes. Using proteomics, we observed a reproducible downregulation of proteins encoded by those mRNAs validated to undergo canonical NGD (Table 1, Figure 6B). We conclude that regulation of mRNA levels by NGD is perpetuated to the protein level. Comparing mRNA to protein abundance (Figure 6A, B), we further observed a subtle but significant trend that proteins ending on strongly affected codons tended to be downregulated, in contrast to those ending on other codons (comparing only proteins translated from mRNAs with >100 CPM in our nanopore sequencing data, to allow comparison with sequencing data itself (strong vs. mild: p = 1.28 × 10^−5^; strong vs. not: p = 1.50 × 10^−4^)). In comparison, it also seems that proteins are generally more strongly downregulated than the mRNAs encoding them (Figure 6A, B). This discrepancy may be explained by additional pathways triggered by ribosome collisions: RQC, which would degrade affected peptides. An involvement of RQC at this point is likely, as the *NEW1* gene also displays positive genetic interactions with the *LTN1* gene (46) (aka *RKR1*; encoding an integral part of the canonical RQC pathway). In conclusion, we showed that New1 protects multiple mRNAs that are characterized by their C-terminal codon, from recruitment of Hel2 and subsequent NGD. This has pervasive consequences on the proteome, which may be, in part, due to additional effects on RQC and possibly other pathways.

### UV crosslinking and analysis of cDNA of reveals New1 interactions with transfer, ribosomal, and messenger RNA

Finally, to find how New1 might exert this activity, we performed CRAC using strains in which New1 was (HTP) tagged, and in which collision sensor Hel2 was either present or absent. We compared these data to complementary CRAC data for Hel2 in presence and absence of New1. Here, we found that New1 binds mRNA, rRNA, and tRNA, consistent with a role in translation (elongation and/or termination) (Table 2).

**Table 2:** Overview of types of RNA that non-collapsed reads were aligned to, comparing New1-CRAC to Hel2-CRAC. New1 binds roughly similar amounts of mRNA and rRNA as Hel2 does, but shows higher binding for tRNA than Hel2. Non-collapsed data are shown here, as collapsing leads to underestimation of highly abundant tRNAs and rRNAs, due to the nature of our unique molecule identifiers.

|  | New1,<br>rep. 1 | New1,<br>rep. 2 | New1,<br>*hel2Δ*,<br>rep. 1 | New1,<br>*hel2Δ*,<br>rep. 2 | Hel2,<br>rep. 1 | Hel2,<br>rep. 2 | Hel2, *new1Δ*,<br>rep. 1 | Hel2, *new1Δ*,<br>rep. 2 |
| --- | --- | --- | --- | --- | --- | --- | --- | --- |
| mRNA | 40.3% | 42.6% | 40.9% | 42.4% | 51.9% | 49.0% | 44.0% | 36.3% |
| rRNA | 35.0% | 34.3% | 41.0% | 31.7% | 36.8% | 39.8% | 43.6% | 53.7% |
| tRNA | 19.7% | 16.4% | 13.5% | 18.6% | 6.6% | 7.4% | 8.0% | 6.9% |
| Other RNAs | 4.9% | 6.6% | 4.5% | 7.3% | 4.6% | 3.7% | 4.4% | 3.1% |

#### New1 interacts with tRNA

The class of New1-bound RNAs that displays the highest difference to Hel2 is tRNA, representing a substantial 13.5 – 19.7% of total (non-collapsed) reads in New1-CRAC libraries. Here, we found tR(CCU), the tRNA that decodes the queue--inducing codon AGG, to be highly bound, representing between 0.5 and 8.3% of total CRAC reads, and between 6.2 and 37.7% of total tRNA reads. Despite the apparent inter--replicate variability, this tRNA species represented one of the most bound tRNA species in all New1 CRAC libraries (Supplementary Table S8). Surprisingly, however, tRNAs that decode other strongly affected codons AAA (tK(UUU)) or CGU (tR(ACG/ICG)), were less represented in the New1 CRAC libraries, compared to some other tRNAs decoding various non-affected codons. Structure-wise, it is unclear how New1 might contact tRNAs, and if this happens while New1 is associated with ribosomes. Considering the flexible N-(140 amino acids missing) and C-termini (84 amino acids missing) that have not been resolved in the published cryo-EM structure (55), it might be possible for them to reach tRNAs within translating or stalled ribosomes. However, our current data do not allow for more detailed conclusions.

#### New1 interacts with the ribosomal small and large subunits and influences ribosomal interactions of collision sensor Hel2

One of the major crosslinked rRNA species was 18S rRNA, in both New1 and Hel2. We used data on single nucleotide micro-deletions, which are frequently induced by nucleotide-amino acid crosslinks (57) to pinpoint sites of direct interaction. In the 18S rRNA, one major (U1361) and two minor (U1491; U493/C495) crosslink sites were observed for New1, where the latter was less prominent. These crosslink sites coincide with the ones previously reported for Hel2 (21), though with a changed preference:

New1 binds more strongly at one of Hel2’s minor binding sites, based on the number of reads mapping to the respective positions (Figure 7A). New1’s major crosslink site, U1361, is one of the minor crosslink sites observed for Hel2, whereas U1491 is Hel2’s major crosslink site. In the published cryo-EM structure of a New1-bound ribosome, this position is in contact with New1. In the absence of Hel2, relative New1 binding at this major crosslink site appeared slightly reduced, whereas binding at the minor crosslink site (U1491; Hel2’s major crosslink site) appeared slightly increased, suggesting that binding of Hel2 at this site, or Hel2-catalyzed poly-ubiquitination of the nearby ribosomal protein Rps20 (21) might hamper New1 binding. Another explanation could be that New1 might bind the minor crosslink site in the context of collided ribosomes, which would probably be more abundant in the absence of Hel2, due to lack of this canonical rescue pathway, leading to an increase in binding at this position. By contrast, Hel2’s binding at its major crosslink site U1491 was markedly increased in the absence of New1, whereas binding at its minor crosslink site U1361 was relatively decreased. This could be due to the increase in ribosome queuing and collisions observed in the absence of New1 (55, 76). It is also possible that *new1Δ*-induced queues represent a different structure from that of standard collided ribosomes, or that other factors bind these complexes, leading to a different binding geometry of Hel2 to those complexes. Additional crosslinks were observed for New1 and Hel2 in 5S, 5.8S and 25S rRNA, however, compared to 18S crosslink sites, they appeared minor (Figure 5A, left). One crosslink site stood out though: a crosslink site at position U33 of 5S rRNA (Figure 5A, right). Although the 5’-half of 5S rRNA is hardly covered by reads, U33 was the second highest deletion site of this RNA species, being deleted in >10% of reads extending beyond this point. This suggests the position to be a prominent interaction site. Like New1’s major 18S rRNA crosslink site, U33 of 5S rRNA clearly makes contact with New1 in the published cryo-EM structure (Figure 5B). This indicates that the structure resolved by cryo-EM, which was determined *ex vivo*, is also relevant in living yeast cells. Our additionally observed 18S rRNA crosslink sites may also be compatible with the cryo-EM structure and could be bound simultaneously, considering that large parts of the New1 N- and C-terminus were not resolved by cryo-EM and are predicted to be highly flexible (Figure 5B). Both, the N- and C-terminus contain positively charged Lys and Arg, as well as aromatic residues, which can mediate RNA binding. As additional evidence for this hypothesis, a crosslinking and mass spectrometry-based technique named identification of RNA-associated peptides (iRAP) (90) determined an RNA binding site at position Arg1170, close to the C-terminus (Figure 5B), which is not visualized by cryo-EM. This position is predicted by alphafold (AF-Q08972-F1) to be in an alpha helix, but is linked to the core of New1 by a long, disordered region. We therefore assume that this position could reach the more distant 18S rRNA crosslink sites.

**Figure 7.**
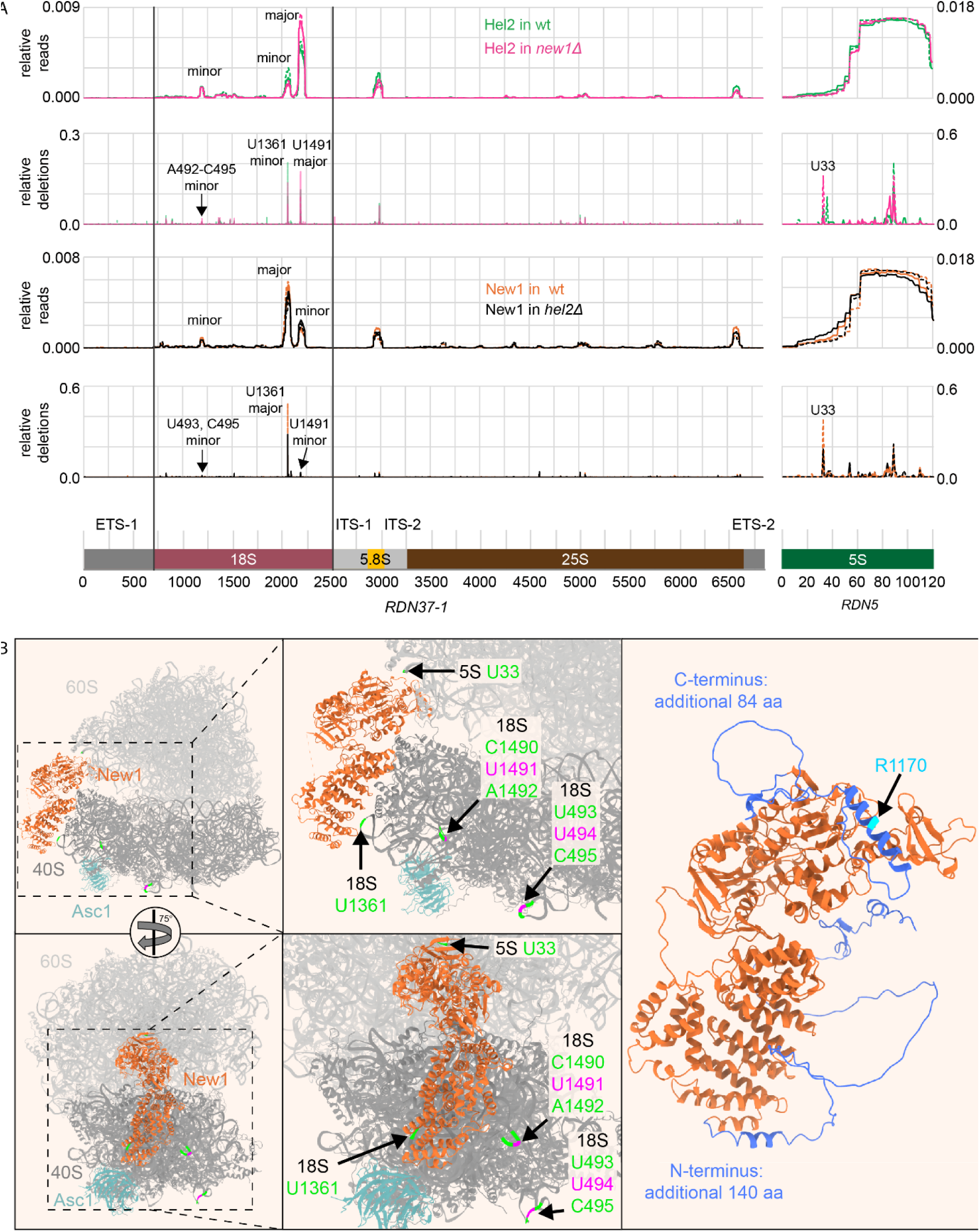
(**A**) CRAC data for Hel2 and New1 interactions with ribosomal RNA at 20°C. Reads and deletions mapped to *RDN37-1* (encoding 18S, 5.8S and 25S rRNAs as a single transcript flanked by external and internal transcribed spacers (ETS, ITS, respectively), as well as *RDN5*, encoding 5S rRNA. For *RDN5*, all reads mapped randomly to either *RDN5-1, RDN5-2, RDN5-3, RDN5-4, RDN5-5* or *RDN5-6* were summed up to compensate for differences arising from random mapping. The sum of reads and deletions mapped to one of the two loci was normalized to 1 and relative reads and relative deletions over the respective locus are shown. Note a difference in scaling between the two loci. Major and minor peaks were determined based on relative reads mapping to the positions. The crosslink positions were determined based on the main deletion peaks mapped in the respective part of the sequence. (**B**) Main deletion peaks mapped to the structure model for New1 bound to a ribosome (pdb: 6S47 (55)) (left and middle panels. Left: full ribosome shown, right: zoom on the New1-bound portion of the ribosome. Colour scheme: green: crosslink sites found in both, Hel2 and New1 CRAC data; Magenta: crosslink sites only found in Hel2 data. Right panel: alphafold (88) model (AF-Q08972-F1) with portion of New1 resolved in the published structure model shown in orange, additional N- and C-terminal amino acids shown in blue. Cyan: Protein-RNA crosslink site within New1 identified by identification of RNA-associated peptides (iRAP) (89).

As an alternative hypothesis, the minor 18S rRNA crosslink sites, but also 5.8S and 25S rRNA crosslink sites we determined could be due to additional binding modes of New1, e.g., during recruitment, potentially in different conformations, which were not captured by cryo-EM (55, 76), so far. The similarities between RNA binding sites of Hel2 and New1 also suggest that binding of these two factors is likely mutually exclusive. Binding of New1 to (stalled or collided) ribosomes might thereby physically block Hel2 from ubiquitinating slowly terminating ribosomes. Its absence, on the other hand, might enable Hel2 activity, thereby terminally stalling such ribosomes, causing or aggravating ribosome queues, and also triggering downstream responses like NGD.

#### New1 interacts with mRNAs with a preference for 3’-termini, independent of the C-terminal codon

The most-bound RNA species for New1 was mRNA (Table 2). Like Hel2, New1 was bound across the mRNA (Supplementary Figure S9), consistent with a general role in translation, either as an active translation elongation factor, like its homologue eEF3 (50), or sampling ribosomes in different translation states, as recently suggested (76), possibly without exhibiting translation elongation factor activity itself. The fact that overexpression of New1 was able to rescue, to some extent, the growth defect caused by repressed transcription of *YEF3* (55) supports a direct role in translation, which can be exerted at least when necessary. In addition to binding across coding sequences, New1 exhibited a major binding peak in 3’-UTRs. This binding pattern is similar to what we found for Hel2 in the absence of New1, on mRNAs with affected C-terminal codons. Given that New1, like Hel2, binds to the 18S rRNA close to the mRNA entry channel, we hypothesize that binding to 3’-UTRs could be due to the presence of New1 at ribosomes upstream of or at Stop codons, possibly facilitated by the protein’s flexible N- and C-termini. However, contrary to Hel2 binding in the absence of New1, New1’s binding to 3’-UTRs did not depend on the identity of the C-terminal codon (Supplementary Figure S9), suggesting New1 is able to associate with terminating ribosomes independently of the C-terminal codon (and tRNA bound within the ribosome). New1 binding to mRNAs was also not altered when comparing CRAC data in presence and absence of Hel2.

In conclusion, our crosslinking data show rRNA interactions of New1 which are in line with published cryo-EM data, as well as additional interactions with rRNA, mRNA and tRNA which have not been described before, and which suggest that New1 is not exclusively engaging terminating ribosomes on queueing-prone mRNA C-termini, but ribosomes at all positions of all mRNAs. This would be in line with structural data showing New1 bound to ribosomes in different translation states (76).

## DISCUSSION

We propose the following order of events upon New1 deficiency (main steps summarized in Figure 8): Lack of New1 leads to inefficient termination at stop codons, dependent on specific C-terminal codons (most prominently: ‘AAA’, ‘AGG’, ‘CGU’), and favours ribosomal collisions. On few mRNAs, containing the ‘UGA’ stop codon followed by the nucleotide ‘C’ (we identified 20 mRNAs in which a strongly affected C-terminal codon is followed by ‘UGAC’), this can lead to stop codon readthrough (76). On all or most of the mRNAs, however, collisions are sensed by Hel2, which ubiquitinates uS10 of affected ribosomes. At least on some mRNAs, the RQT complex can then split the leading (stalled) ribosome, enabling Cue2-mediated endonucleolytic cleavage within the collided ribosome. Here it is unclear (as it is in general in the current literature) whether that ribosome is first split by the RQT complex, or whether cleavage occurs prior to splitting, which might then more likely be catalyzed by Dom34. Split 60S:peptide complexes are likely targeted to RQC (not further investigated in our study). Alternatively, supported by part of our data, Cue2 may also cleave mRNA upstream of ribosomes, without prior ribosome splitting via the RQC-uncoupled NGD pathway. Endonucleolytic fragments are then degraded exonucleolytically by the exosome with the help of the Ski complex, and by Xrn1 (as is consensus in the literature). The degradation of affected mRNAs further propagates to the level of corresponding proteins, where several essential metabolic enzymes (e.g., Pgk1 and Gpm1) and translation components, which are among the most abundant yeast proteins (e.g., eEF-1 alpha and eEF-1 beta) are markedly reduced. Noteworthy, most mRNAs were not covered sufficiently in our Nanopore-Seq experiment to assess whether or not they had been cleaved by Cue2.

**Figure 8:**
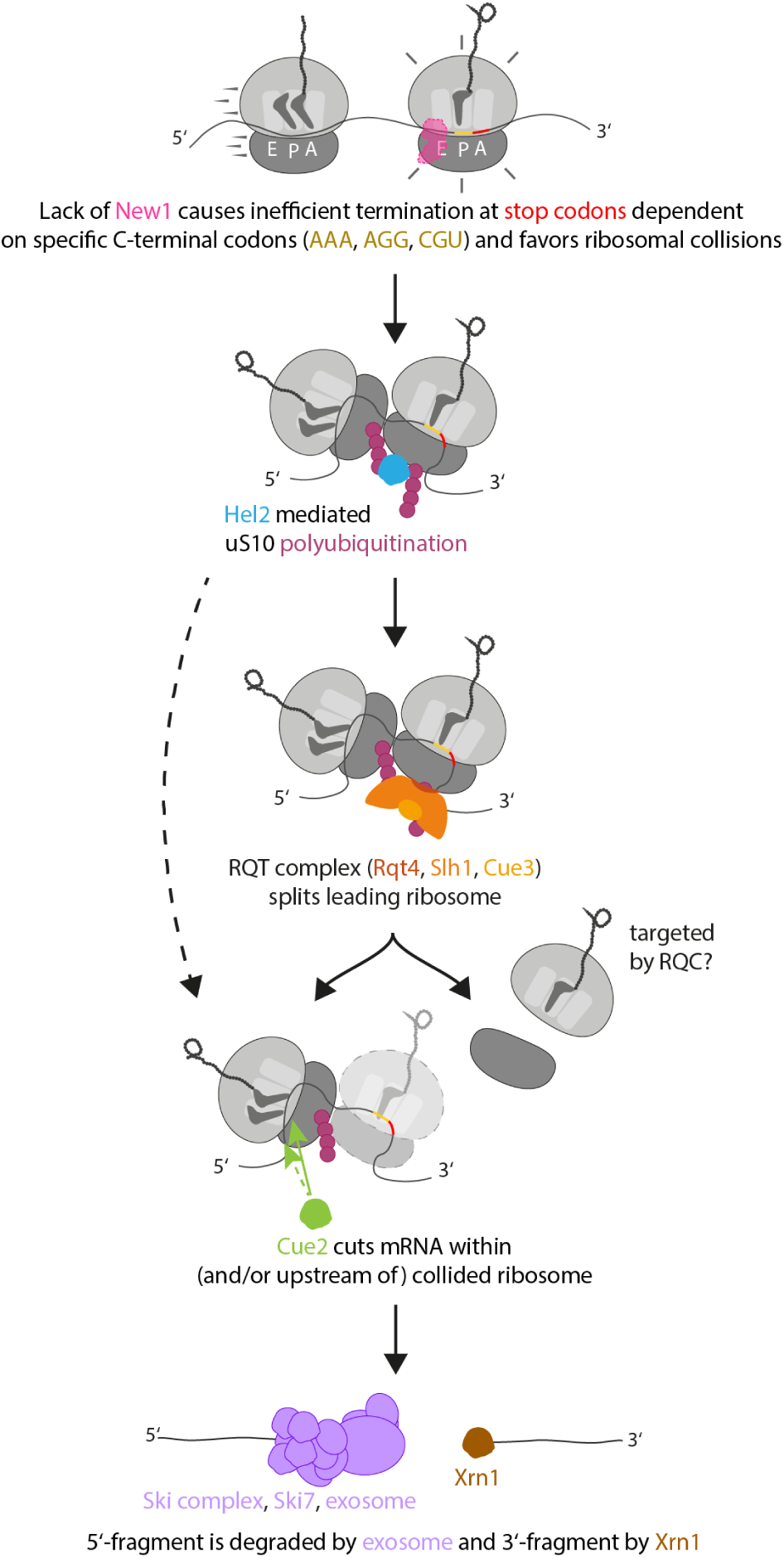
Proposed order of events in the absence of New1. Dashed arrows represent RQC-uncoupled NGD, where Cue2 cleaves at the 5’-end of the collision, without the need for RQT-catalyzed splitting. The translucent ribosome would only be present in the RQC-uncoupled pathway.

However, we did observe examples that were clearly not targets of Cue2, although queueing and Hel2 recruitment happen for them (Rps15, Rpl41B). What governs the decision of whether or not an mRNA is targeted to NGD upon Hel2 recognition remains to be elucidated. It is possible that, on some mRNAs, the RQT can split ribosomes to either allow for translation to resume, or to target peptides to decay via RQC, without eliciting NGD. In addition, we observed that the relative decrease in protein levels translated from strongly affected mRNAs was greater than the decrease in mRNA levels, many of which remained unchanged (Figure 6). Here as well, it is possible that the decrease in protein levels is more strongly influenced by the RQC pathway, which can occur in parallel to or independently of NGD. Although we did not investigate the effect of New1 on RQC in this work, we note that the *NEW1* gene exhibits positive genetic interactions with the *RKR1/LTN1* gene, which encodes the central RQC factor Ltn1, responsible for ubiquitination of nascent peptides on stalled ribosomes. We therefore find it highly likely that RQC-based degradation of peptides from collided ribosomes is also triggered in the absence of New1.

Although the growth defect upon deletion of *NEW1* is more pronounced at 20°C than 30°C, ribosome collisions (55, 76), Hel2 recruitment and downregulation of mRNA appeared to happen to a similar extent according to recently published (76), as well as our data (Figure 1, Table 1). One possible explanation for this could be that one (or several) of the downregulated genes is more important for growth at 20°C than 30°C, or that one or few factors are more affected at 20°C than 30°C. This would be possible, as single mRNAs appeared to be affected to varying degrees at both temperatures (Table 1).

Interestingly, we observe an over-representation of strongly affected lysine and arginine codons at C-termini, compared to their frequency in gene bodies in *S. cerevisiae*, whereas alternative, non-affected codons encoding the same amino acid are under-represented at C-termini (Table 3). This is despite the fact that several important cellular proteins are, as a consequence, strongly reduced upon lack of New1. This striking over-representation may suggest a potential fitness benefit by retaining, or even favouring queue-inducing C-terminal codons over iso-coding non-queue-inducing C-terminal codons (e.g. favouring K(AAA) over K(AAG)), when New1 is available. The over-representation of queuing-prone C-terminal codons, which requires New1 to avoid fitness losses also suggests that New1 could have regulatory potential. As an example, New1 expression was found to be strongly decreased in stationary phase (91), where this may contribute to downregulation of New1’s target genes. As another hypothesis, the highly affected C-terminal codons may allow longer time for termination, e.g., to allow proper folding or binding of co-factors/interactors, before peptide release, while still maintaining proper termination in the presence of New1. Although we are beginning to better understand the consequences of lack of New1, what exactly the protein does is still unclear. Cryo-EM studies (55, 76) suggest that its binding mode is similar to that of its better characterized homologue eEF3. Our crosslinking data also support that New1 interacts with ribosomes *in vivo*, and that it does so along the whole mRNA, however with a preference for the 3’-terminus, suggesting a preference for terminating ribosomes. Although the absence of New1 only induces ribosome collisions and related downstream consequences on specific mRNAs, New1 itself is apparently able to interact with terminating ribosomes on any mRNA. So why does lack of New1 not affect all mRNAs and terminating ribosomes equally? We suggest that another, additional factor, e.g., eEF3, which has also been implicated in translation termination (53), is able to fulfil the same role as New1 on most, but not all mRNAs/tRNAs. This would be in line with published findings that New1 overexpression was able to compensate to some degree for eEF3 deficiency, but presence of eEF3 was not able to compensate for *NEW1* deletion (55). Therefore, specifically in the context of translation termination, New1 seems to be required only for a subset of mRNAs/tRNAs. On the structural level, this differential requirement could be caused by the fact that the chromodomain, which, at least in the case of eEF3, interacts with the ribosome’s L1 stalk (50) is shorter for New1 than for eEF3 (55). Alternatively, New1 could be probing all terminating ribosomes, but only act on a subset of them that exhibits problems in termination. Such problems could be caused, e.g., by the local structure of mRNA, tRNA and/or termination factors within the ribosome. Indeed, it is highly likely that the tRNAs that decode affected codons are major determinants of the problems in termination and the queueing phenotype. tRNA specificity probably explains the codon dependence we observe. It has been convincingly demonstrated that deletion of the main tRNA decoding the ‘AGG’ codon (HSX1/tR(CCU)) inhibited C-terminal ribosome queueing upon deletion of *NEW1* (76). In addition, specific tRNAs have very recently been shown to recruit the CCR4-NOT complex to human translating ribosomes, triggering mRNA decay, whereas structural properties of other tRNAs prevent this recruitment (92). In a similar fashion, tRNA structure could hamper proper recruitment of termination factors to the terminating ribosome, as suggested by Turnbull et al. (76). As an additional feature, tRNA modification could be a determinant of termination problems (76). However, arginine codons R(CGA) and R(CGC) and R(CGU) are all decoded by the same tRNA, but are not affected by lack of New1 to the same extent (CGA, CGC – mildly affected; CGU – strongly affected). This suggests that it is not the tRNA alone that determines the extent of problems in termination. A role for mRNA structure has previously been demonstrated for difficult-to-decode codon stretches, where the conformation of mRNA within the ribosome interferes with tRNA binding (8, 9). Finally, we could also envision that the local structure of the ribosome itself could play a role, where, e.g., the ribosomal L1 stalk might be in different conformations, depending on the P/E-site or E-site tRNA.

**Table 3:**
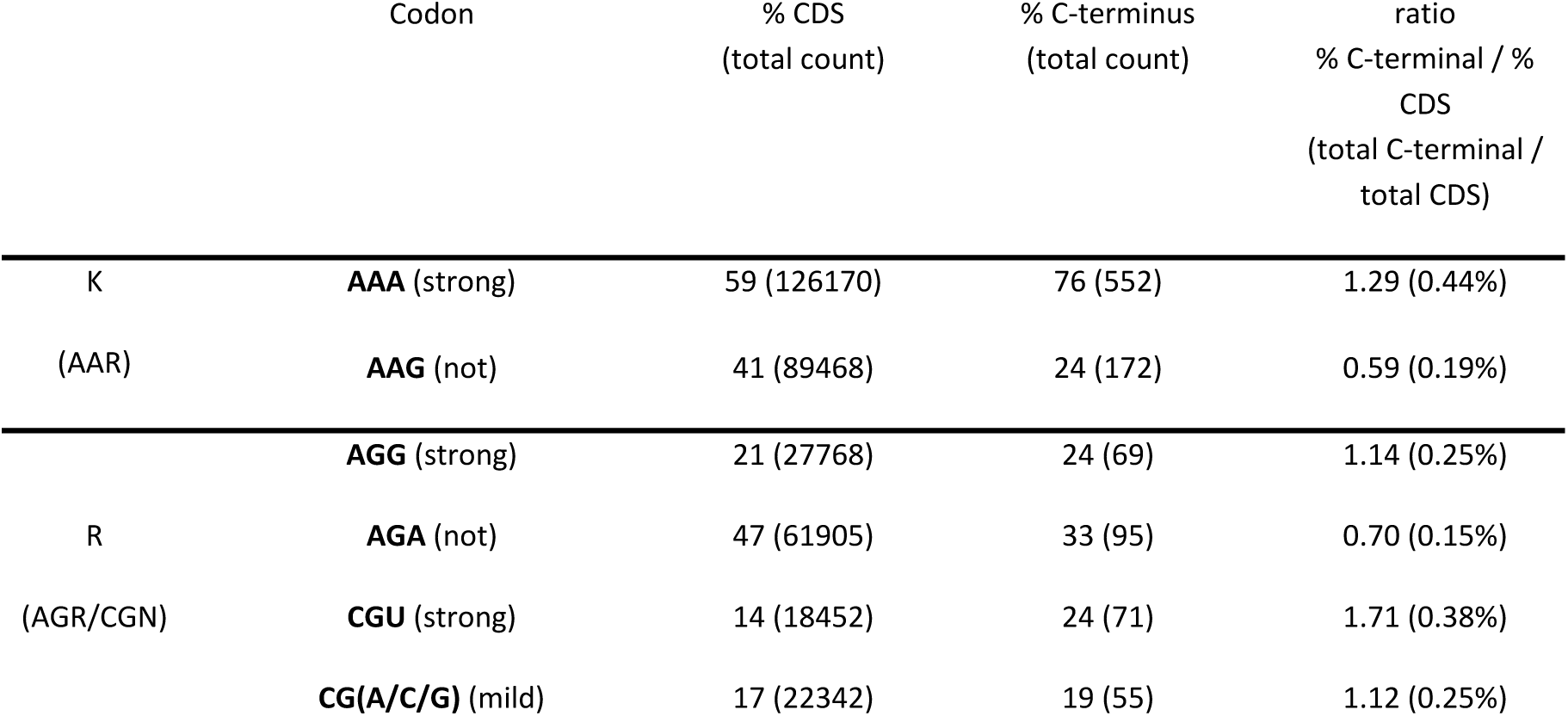
C-terminally stalling codons are overrepresented at the C-terminus in *S. cerevisiae* (from our own reference files, see Materials and Methods). Percentage of specific lysine and arginine codons of total lysine or arginine codons at C-termini and in CDS are given, as well as ratio of percentage at C-terminus and CDS.

Although ribosome collision at the stop codon is an important phenotype in *new1Δ*, the structure of the collided ribosomes from such strains has not been determined yet. Since these collisions initiate with ribosomes containing the stop codon in the A-site, it remains to be seen whether these ribosomes already contain termination factors, or whether the A-site remains vacant, e.g., due to structure of the P-site tRNA. At least in presence of New1, ribosomes containing both, New1 and eRF1 have been observed (76). Such variations may differentiate C-terminal collided ribosomes from ribosomes collided on other stall-inducing sequences, and may also have consequences on downstream surveillance pathways.

Beyond giving insights into the function, as well as consequences of dysfunction of New1, our work exemplifies, to our knowledge for the first time, how (in the absence of New1) highly abundant proteins are mRNA-specifically downregulated by the NGD pathway, under physiological conditions, in the absence of translation inhibitors or mRNA damage. New1 represents a protective factor that is needed to avoid this form of degradation.

## DATA AVAILABILITY

CRAC data are available at GEO, under the accession number GSE275093. Nanopore sequencing data are available at Bioproject under the BioSample accessions <u>SAMN43836079</u> –<u>SAMN43836090</u> (Project Number: PRJNA1159148). The mass spectrometry proteomics data have been deposited to the ProteomeXchange Consortium via the PRIDE (93) partner repository with the dataset identifier <u>PXD055983</u>.

## SUPPLEMENTARY DATA

Supplementary Data are available.

## AUTHOR CONTRIBUTIONS

M.L.W. conceived the study; M.L.W. and M.M. designed experiments; M.M. performed and analyzed all biochemical experiments, including Western and Northern blots; M.L.W., L.S.T and E.P. performed CRAC experiments; and analysis, M.L.W., L.S.T. and M.M. analyzed CRAC experiments; M.M. and T.B. performed nanopore library preparation and sequencing, S.P., T.B., M.M., L.S.T. and M.L.W. analyzed nanopore sequencing data, M.M., L.S.T. and M.L.W. re-analyzed published riboSeq data; M.M. and M.L.W. analyzed proteomics data; M.M. and K.I. re-analyzed published cryoEM data. All authors wrote the manuscript and discussed the data.

## ACKNOWLEDGEMENTS

We gratefully acknowledge David Tollervey (University of Edinburgh) for his generous support of our CRAC experiments. We acknowledge the Proteomics Core Facility at Institute of Molecular Biology (IMB) Mainz for proteomics analysis including sample processing, mass spectrometry measurement and data analysis as well as data upload to the ProteomeXchange Consortium. We thank Susanne Gerber (Johannes Gutenberg-University Mainz) for kindly allowing us to use the PromethION device as part of our study. We further acknowledge Hans Zischler and Maurice Scheuren (Johannes Gutenberg-University Mainz) for reagents and the loan of a hybridisation oven, Praveen Bawankar (Johannes Gutenberg-University Mainz) for support with the establishment of Northern blotting, Kristina Friedland (Johannes Gutenberg-University Mainz) for access to the FUSION PULSE TS gel documentation system and Mark Helm (Johannes Gutenberg-University Mainz) for access to equipment for quantification of nucleic acids. We further thank Kamena Kostova (Stower’s Institute for Medical Research) for plasmid pKK148, Helle Ulrich (Institute of Molecular Biology; Johannes Gutenberg-University Mainz) for plasmid YEp181-CUP1-His-Ub, and Brian Luke (Johannes Gutenberg-University Mainz) for plasmids pYM18 and pYM20. Finally, we also thank Marcel Hüttel (Johannes Gutenberg-University Mainz) for the 3D print of an agarose gel spacer.

