## Supplementary Data File for "Yeast elongation factor homolog New1 protects a subset of mRNAs from degradation by no-go decay"

### Content:

**Supplementary Figure S1** **p. 2**

**Supplementary Figure S2** **p. 3**

**Supplementary Figure S3** **p. 3**

**Supplementary Figure S4** **p. 4**

**Supplementary Figure S5** **p. 5**

**Supplementary Figure S6** **p. 6**

**Supplementary Figure S7** **p. 6**

**Supplementary Figure S8** **p. 7**

**Supplementary Figure S9** **p. 8**

**Supplementary Table S1** **p. 9**

**Supplementary Table S2** **p. 12**

**Supplementary Table S3** **p. 15**

**Supplementary Table S4** **p. 16**

**Supplementary Table S5** **p. 19**

**Supplementary Table S6** **p. 20**

**Supplementary Table S7** **p. 21**

**Supplementary Table S8** **p. 22**

**Supplementary References** **p. 23**

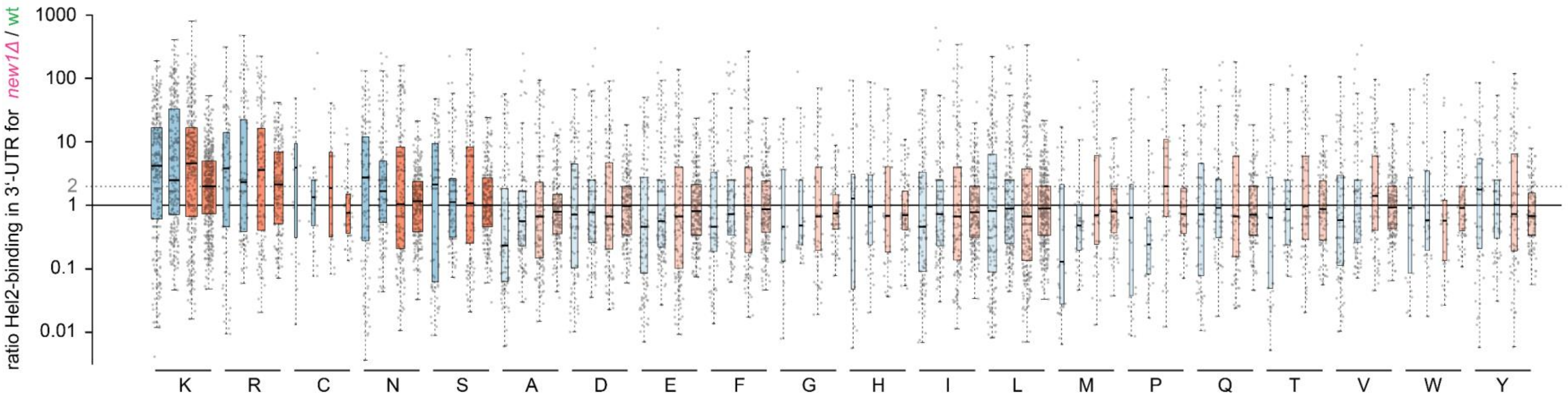


**Supplementary Figure S1.** Ratio of Hel2 binding in 3′-UTR for *new1Δ*/wt at 20°C (blue, 2 replicates), and 30°C (red, 2 replicates) for mRNAs, plotted by C-terminally encoded amino acids. Non-affected amino acids are shown with higher transparency. Center lines show the medians, box limits indicate the 25$^{th}$ and 75$^{th}$ percentiles, as determined by R software; whiskers extend 1.5 times the interquartile range from the 25$^{th}$ and 75$^{th}$ percentiles. All data points are represented by dots. Width of the boxes is proportional to the square root of the sample size. Related to **Figure 2**.

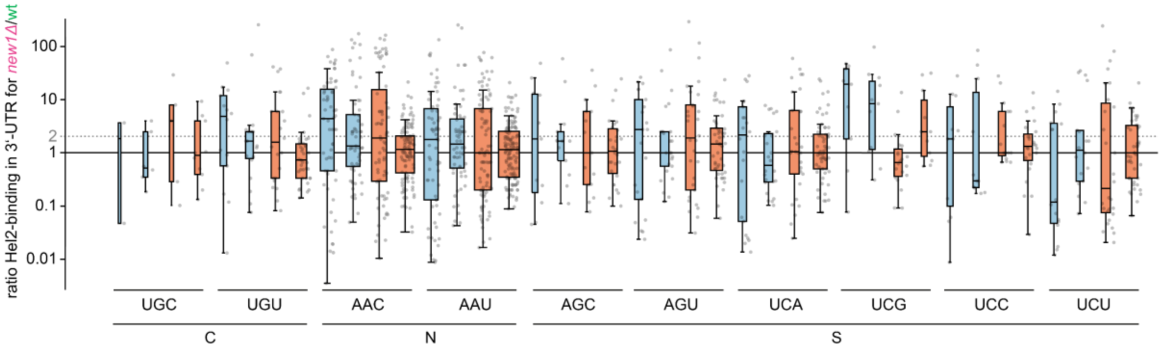


**Supplementary Figure S2.** Ratio of Hel2 binding in 3'-UTR for *new1Δ*/wt for all cysteine, asparagine, and serine codons at 20°C (blue, 2 replicates), and 30°C (red, 2 replicates). Center lines show the medians, box limits indicate the 25th and 75th percentiles, as determined by R software; whiskers extend 1.5 times the interquartile range from the 25th and 75th percentiles. All data points are represented by dots. Width of the boxes is proportional to the square root of the sample size. Related to **Figure 2.**

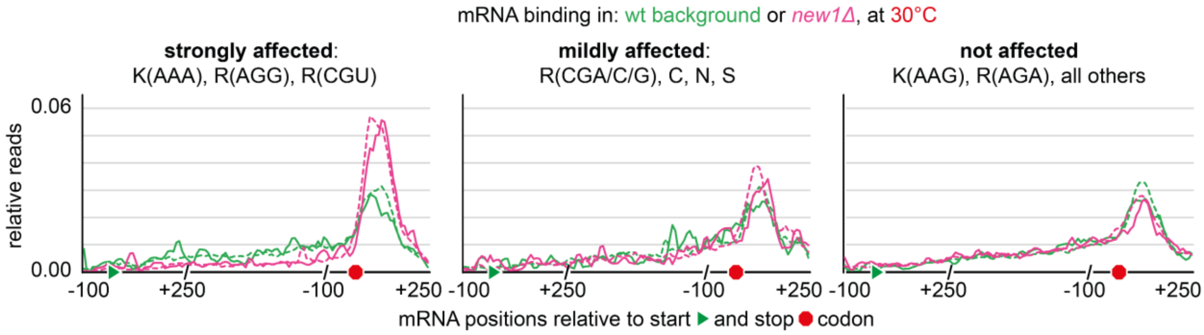


**Supplementary Figures S3.** Metaplots showing Hel2 binding (relative hits) in either wildtype (wt, green, 2 biological replicates) or *new1Δ* (magenta, 2 biological replicates) background, at 30°C, for those mRNAs within the group of top 1000 highest Hel2-bound mRNAs (1) with C-terminal codons being either strongly affected, mildly affected or not affected by lack of New1. **Related to Figure 2**.

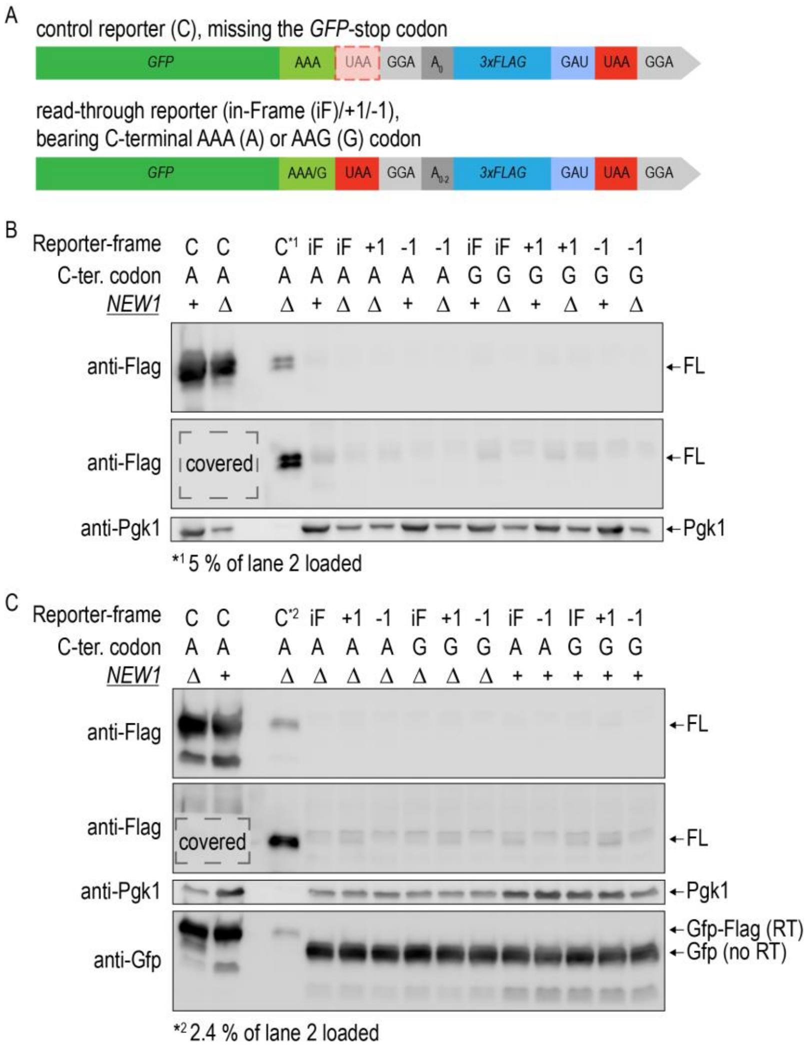


**Supplementary Figure S4.** Readthrough reporter assay. (**A**) The readthrough reporter consist of the coding sequence of GFP, with either the native lysine codon 'AAA', found to induce queuing upon *new1Δ,* or the non-queueing C-terminal lysine codon 'AAG', followed by the strong stop codon 'UAA'. To avoid sequence variation downstream of the stop codon, a glycine spacer ('GGA') was added. To allow tagging of readthrough products with a 3xFLAG-tag in all three reading frames, 3 constructs were designed, containing either no, one or two additional nucleotides 'A' for in-frame, +1 or -1 reading frame, respectively. Since the 3xFLAG sequence would end with a lysine codon ('AAG'), an additional aspartic acid ('GAU') spacer was introduced in front of the 3xFLAG stop codon (UAA), followed by a glycine codon ('GGA'). The control construct is identical to the described reporters, except that it misses the GFP stop codon. Consequently, each ribosome is capable of carrying out translation to the end of the 3xFLAG-tag. (**B**) Western blot analysis of GFP-3xFLAG-tagged readthrough reporter protein in *new1Δ* (*NEW1*: Δ) and wt (*NEW1*: +) strains harbouring either the control construct ('C') or constructs with the 3xFLAG-tag in-frame (iF), in the +1 shifted frame (+1) or in the -1 shifted frame (-1), with a C-terminal lysine codon 'AAA' ('A') or 'AAG' ('G'). Protein extract from equal amounts of cells was loaded per lane. In the second panel, the control samples were covered during visualization. Indices show the relative loading compared to lane 2 (**B**) or lane 1 (**C**). In (**C**) the Gfp signal is visualized in addition and sizes of Gfp-Flag with readthrough (RT) or without readthrough (no-RT) are indicated.

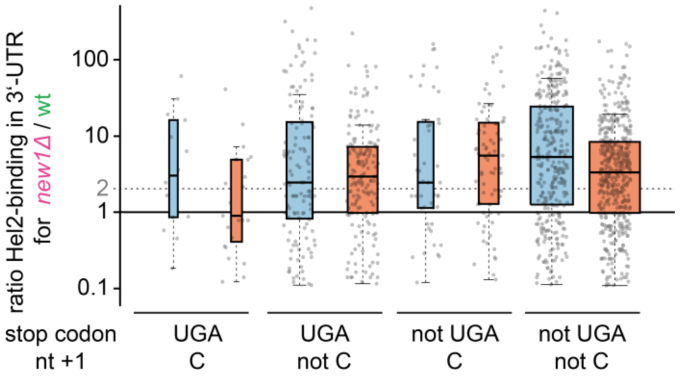


**Supplementary Figure S5.** Boxplot analysis of average Hel2 3′-UTR enrichment for mRNAs, containing strongly affected C-terminal codons at 20°C (blue, data from 2 biological replicates combined) and 30°C (red, data from 2 biological replicates combined), depending on stop codon context. mRNAs were sorted into the following groups: (1) stop codon 'UGA', followed by nucleotide 'C' at position +1; (2) stop codon 'UGA' and a nucleotide 'A', 'G', or 'U' a position +1, (3) stop codon 'UAA' or 'UAG', followed by nucleotide 'C' at position +1; (4) stop codon 'UAA' or 'UAG' and a nucleotide 'A', 'G', or 'U' a position +1. Center lines show the medians, box limits indicate the 25th and 75th percentiles, as determined by R software; whiskers extend 1.5 times the interquartile range from the 25th and 75th percentiles. All data points are represented by dots. Width of the boxes is proportional to the square root of the sample size.

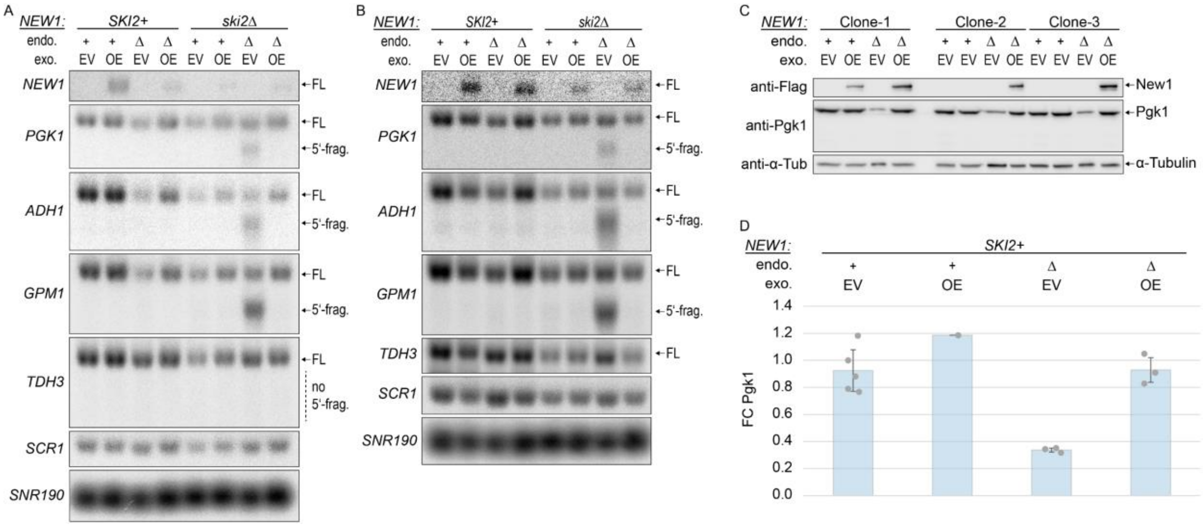


**Supplementary Figure S6.** Further replicates for study of NGD fragments at 20°C. Related to **Figure 3.** (**A, B**) Northern blot analysis of RNA levels for *NEW1*, *PGK1*, *ADH1*, *GPM1*, *TDH3*, *SCR1*, and *SNR190* in *new1Δ* (endo Δ), *NEW1*-positive (endo +) strains, containing an empty vector (exo EV) or New1-overexpression vector (exo OE), and containing or lacking the *SKI2* gene. Blots for replicates 2 and 3 (quantified in **Figure 3B**). (**C**) Western blot analysis of Pgk1 levels in *new1Δ* (endo Δ; exo EV), wt-like (endo +; exo EV), and New1-overexpressing (endo+ or endo Δ; exo OE) strains, compared to wildtype strains containing an empty vector (endo +, exo EV). (**D**) Quantification of protein levels for Pgk1, relative to Tubulin, related to (**C**), n = 3 biological replicates for endo Δ; exo EV and endo Δ; exo OE, n = 5 biological replicates for endo + ; exo EV, n = 1 biological replicates for endo +; exo OE. All data quantified as FC compared to median of all endo + values. Error bars represent 1 standard deviation.

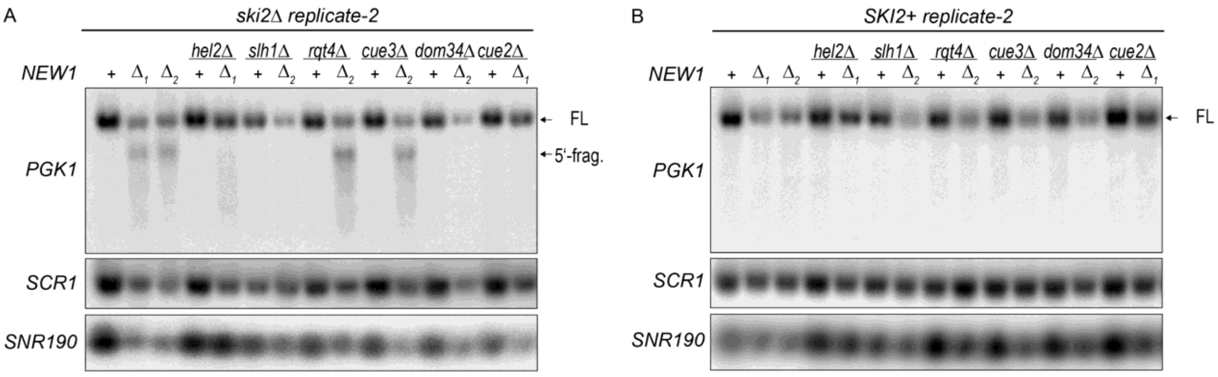


**Supplementary Figure S7.** Further replicates for study of NGD fragments in mutants lacking additional NGD players, at 20°C. Related to **Figure 4**. Additional biological replicate for Northern blot analysis of no-go decay dependence on various NGD factors. (**A**) in *ski2Δ* strains, (**B**) in *SKI2* containing strains. Quantification in **Figure 4**. $\Delta_1$ designates mutants in which the *NEW1* gene was replaced with a KanMX cassette, $\Delta_2$ designates mutants in which the *NEW1* gene was replaced with a HphMX cassette. FL: full-length; 5′-frag.: endonucleolytic 5′-fragment.

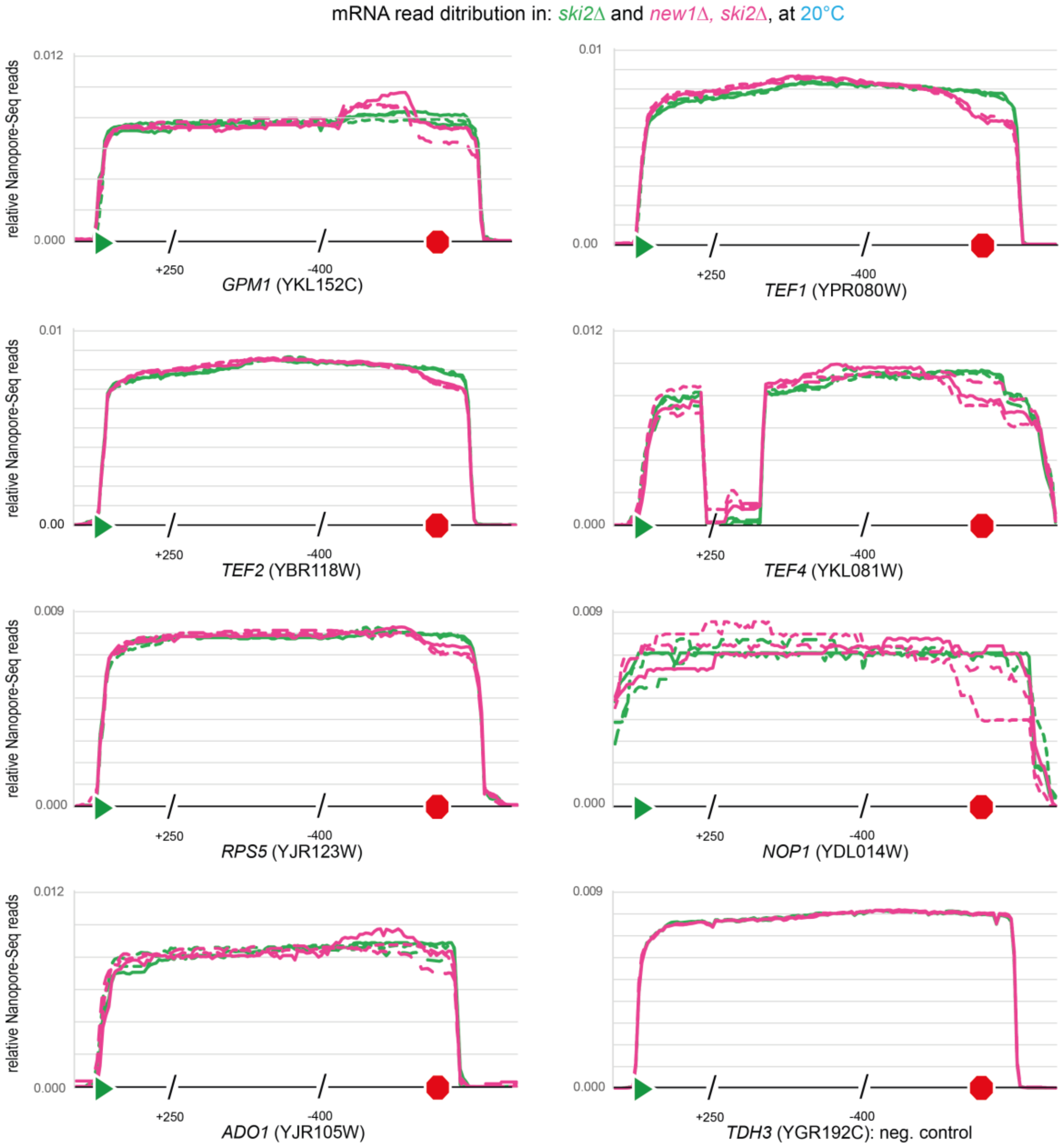


**Supplementary Figure S8.** Analysis of endonucleolytic fragments via Nanopore-Seq in *new1Δ, ski2Δ* double knockout and *ski2Δ* single knockout yeast strains. Relative Nanopore-Seq read distribution along the *GPM1*, TEF*1, TEF2, TEF4* (including an intron), *RPS5, NOP1, and ADO1* transcripts, which end on strongly queueing-prone C-terminal codons. The *TDH3* transcript (ending on non-affected codon 'GCU', encoding alanine. 100 nt upstream of and 250 nt downstream of the start codon, as well as 400 nt upstream of and 250 nt downstream of the stop codon are shown unscaled, whereas the rest of the gene body was scaled. Related to **Figure 5**.

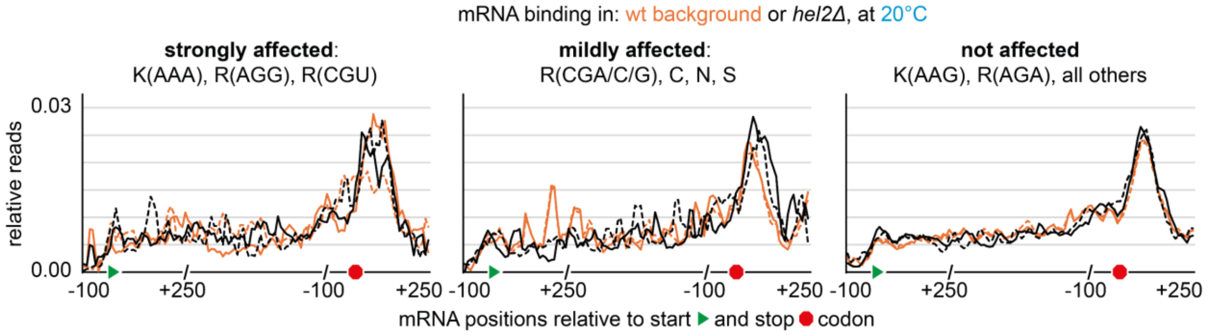


**Supplementary Figure S9.** Metaplots showing New1 binding (relative reads) in either wildtype (wt, orange, 2 biological replicates) or *hel2Δ* (black, 2 biological replicates) background, at 20°C, for those mRNAs within the group of top 1000 highest Hel2-bound mRNAs (1) with C-terminal codons being either strongly affected, mildly affected or not affected by lack of New1.

**Supplementary Table S1.** List of strains used in this study

| # | Strain | Genotype | Parental strain | Reference or source |
|---|---|---|---|---|
| 1 | BY4741 | MATa, his3Δ, *LEU2Δ*, met15Δ, ura3Δ | - | Ref. (2) |
| 2 | *new1Δ*::KanMX | BY4741, *new1Δ*::KanMX | BY4741 (1) | this study |
| 3 | *new1Δ*::HphMX | BY4741, *new1Δ*::HphMX | BY4741 (1) | this study |
| 4 | *hel2Δ* | BY4741, *hel2Δ* | BY4741 (1) | Ref. (1) |
| 5 | *hel2Δ*, *new1Δ*::KanMX | BY4741, *hel2Δ*, *new1Δ*::KanMX | *hel2Δ* (4) | this study |
| 6 | *slh1Δ*::KanMX | BY4741, *slh1Δ*::KanMX | BY4741 (1) | this study |
| 7 | *slh1Δ*::KanMX, *new1Δ*::HphMX | BY4741, *slh1Δ*::KanMX, *new1Δ*::HphMX | *slh1Δ*::KanMX (6) | this study |
| 8 | *rqt4Δ*::KanMX | BY4741, *rqt4Δ*::KanMX | BY4741 (1) | this study |
| 9 | *rqt4Δ*::KanMX, *new1Δ*::HphMX | BY4741, *rqt4Δ*::KanMX, *new1Δ*::HphMX | *rqt4Δ*::KanMX (8) | this study |
| 10 | *cue3Δ*::KanMX | BY4741, *cue3Δ*::KanMX | BY4741 (1) | this study |
| 11 | *cue3Δ*::KanMX, *new1Δ*::HphMX | BY4741, *cue3Δ*::KanMX, *new1Δ*::HphMX | *cue3Δ*::KanMX (10) | this study |
| 12 | *cue2Δ*::KanMX | BY4741, *cue2Δ*::KanMX | BY4741 (1) | this study |
| 13 | *new1Δ*::KanMX, *cue2Δ*::HphMX | BY4741, *new1Δ*::KanMX, *cue2Δ*::HphMX | *new1Δ*::KanMX (2) | this study |
| 14 | *dom34Δ*::KanMX | BY4741, *dom34Δ*::KanMX | BY4741 (1) | this study |
| 15 | *new1Δ*::HphMX, *dom34Δ*::KanMX | BY4741, *new1Δ*::HphMX, *dom34Δ*::KanMX | *new1Δ*::HphMX (3) | this study |
| 16 | *ski2Δ*::*LEU2* | BY4741, *ski2Δ*::*LEU2* | BY4741 (1) | this study |
| 17 | *new1Δ*::KanMX, *ski2Δ*::*LEU2* | BY4741, *new1Δ*::KanMX, *ski2Δ*::*LEU2* | *new1Δ*::KanMX (2) | this study |
| 18 | *new1Δ*::HphMX, *ski2Δ*::*LEU2* | BY4741, *new1Δ*::HphMX, *ski2Δ*::*LEU2* | *new1Δ*::HphMX (3) | this study |
| 19 | *hel2Δ*, *ski2Δ*::*LEU2* | BY4741, *hel2Δ*, *ski2Δ*::*LEU2* | *hel2Δ* (4) | this study |
| 20 | *hel2Δ*, *new1Δ*::KanMX, *ski2Δ*::*LEU2* | BY4741, *hel2Δ*, *new1Δ*::KanMX, *ski2Δ*::*LEU2* | *hel2Δ*, *new1Δ*::KanMX (5) | this study |
| 21 | *slh1Δ*::KanMX, *ski2Δ*::*LEU2* | BY4741, *slh1Δ*::KanMX, *ski2Δ*::*LEU2* | *slh1Δ*::KanMX (6) | this study |
| 22 | *slh1Δ*::KanMX, *new1Δ*::HphMX, *ski2Δ*::*LEU2* | BY4741, *slh1Δ*::KanMX, *new1Δ*::HphMX, *ski2Δ*::*LEU2* | *slh1Δ*::KanMX, *new1Δ*::HphMX (7) | this study |
| 23 | *rqt4Δ*::KanMX, *ski2Δ*::*LEU2* | BY4741, *rqt4Δ*::KanMX, *ski2Δ*::*LEU2* | *rqt4Δ*::KanMX (8) | this study |
| 24 | *rqt4Δ*::KanMX, *new1Δ*::HphMX, *ski2Δ*::*LEU2* | BY4741, *rqt4Δ*::KanMX, *new1Δ*::HphMX, *ski2Δ*::*LEU2* | *rqt4Δ*::KanMX, *new1Δ*::HphMX (9) | this study |
| 25 | *cue3Δ*::KanMX, *ski2Δ*::*LEU2* | BY4741, *cue3Δ*::KanMX, *ski2Δ*::*LEU2* | *cue3Δ*::KanMX (10) | this study |
| 26 | *cue3Δ*::KanMX, *new1Δ*::HphMX, *ski2Δ*::*LEU2* | BY4741, *cue3Δ*::KanMX, *new1Δ*::HphMX, *ski2Δ*::*LEU2* | *cue3Δ*::KanMX, *new1Δ*::HphMX (11) | this study |
| 27 | *cue2Δ*::KanMX, *ski2Δ*::*LEU2* | BY4741, *cue2Δ*::KanMX, *ski2Δ*::*LEU2* | *cue2Δ*::KanMX (12) | this study |
| 28 | *new1Δ*::KanMX, *Cue2Δ*::HphMX, *ski2Δ*::*LEU2* | BY4741, *new1Δ*::KanMX, *Cue2Δ*::HphMX, *ski2Δ*::*LEU2* | *new1Δ*::KanMX, *cue2Δ*::HphMX (13) | this study |
| 29 | *dom34Δ*::KanMX, *ski2Δ*::*LEU2* | BY4741, *dom34Δ*::KanMX, *ski2Δ*::*LEU2* | *dom34Δ*::KanMX (14) | this study |
| 30 | *new1Δ*::HphMX, *dom34Δ*::KanMX, *ski2Δ*::*LEU2* | BY4741, *new1Δ*::HphMX, *dom34Δ*::KanMX, *ski2Δ*::*LEU2* | *new1Δ*::HphMX, *dom34Δ*::KanMX (15) | this study |
| 31 | BY4741, pGTSTR | BY4741, BY4741, pGTSTR | BY4741 (1) | this study |
| 32 | *new1Δ*::KanMX, pGTSTR | BY4741, *new1Δ*::KanMX, pGTSTR | *new1Δ*::KanMX (2) | this study |
| 33 | *new1Δ*::HphMX, pGTSTR | BY4741, *new1Δ*::HphMX, pGTSTR | *new1Δ*::HphMX (3) | this study |
| 34 | *hel2Δ*, pGTSTR | BY4741, *hel2Δ*, pGTSTR | *hel2Δ* (4) | this study |
| 35 | *hel2Δ*, *new1Δ*::KanMX, pGTSTR | BY4741, *hel2Δ*, *new1Δ*::KanMX, pGTSTR | *hel2Δ*, *new1Δ*::KanMX (5) | this study |
| 36 | *slh1Δ*::KanMX, pGTSTR | BY4741, *slh1Δ*::KanMX, pGTSTR | *slh1Δ*::KanMX (6) | this study |
| 37 | *slh1Δ*::KanMX, *new1Δ*::HphMX, pGTSTR | BY4741, *slh1Δ*::KanMX, *new1Δ*::HphMX, pGTSTR | *slh1Δ*::KanMX, *new1Δ*::HphMX (7) | this study |
| 38 | *rqt4Δ*::KanMX, pGTSTR | BY4741, *rqt4Δ*::KanMX, pGTSTR | *rqt4Δ*::KanMX (8) | this study |

| | | | | |
|---|---|---|---|---|
| 39 | *rqt4Δ*::KanMX, *new1Δ*::HphMX, pGTSTR | BY4741, *rqt4Δ*::KanMX, *new1Δ*::HphMX, pGTSTR | *rqt4Δ*::KanMX, *new1Δ*::HphMX (9) | this study |
| 40 | *cue3Δ*::KanMX, pGTSTR | BY4741, *cue3Δ*::KanMX, pGTSTR | *cue3Δ*::KanMX (10) | this study |
| 41 | *cue3Δ*::KanMX, *new1Δ*::HphMX, pGTSTR | BY4741, *cue3Δ*::KanMX, *new1Δ*::HphMX, pGTSTR | *cue3Δ*::KanMX, *new1Δ*::HphMX (11) | this study |
| 42 | *dom34Δ*::KanMX, pGTSTR | BY4741, *dom34Δ*::KanMX, pGTSTR | *dom34Δ*::KanMX (14) | this study |
| 43 | *dom34Δ*::KanMX, *new1Δ*::HphMX, pGTSTR | BY4741, *dom34Δ*::KanMX, *new1Δ*::HphMX, pGTSTR | *new1Δ*::HphMX, *dom34Δ*::KanMX (15) | this study |
| 44 | *cue2Δ*::KanMX, pGTSTR | BY4741, *cue2Δ*::KanMX, pGTSTR | *cue2Δ*::KanMX (12) | this study |
| 45 | *new1Δ*::KanMX, *cue2Δ*::HphMX, pGTSTR | BY4741, *new1Δ*::KanMX, *cue2Δ*::HphMX, pGTSTR | *new1Δ*::KanMX, *cue2Δ*::HphMX (13) | this study |
| 46 | *ski2Δ*::*LEU2*, pGTSTR | BY4741, *ski2Δ*::*LEU2*, pGTSTR | *ski2Δ*::*LEU2* (16) | this study |
| 47 | *new1Δ*::KanMX, *ski2Δ*::*LEU2*, pGTSTR | BY4741, *new1Δ*::KanMX, *ski2Δ*::*LEU2*, pGTSTR | *new1Δ*::KanMX, *ski2Δ*::*LEU2* (17) | this study |
| 48 | *new1Δ*::HphMX, *ski2Δ*::*LEU2*, pGTSTR | BY4741, *new1Δ*::HphMX, *ski2Δ*::*LEU2*, pGTSTR | *new1Δ*::HphMX, *ski2Δ*::*LEU2* (18) | this study |
| 49 | *hel2Δ*, *ski2Δ*::*LEU2*, pGTSTR | BY4741, *hel2Δ*, *ski2Δ*::*LEU2*, pGTSTR | *hel2Δ*, *ski2Δ*::*LEU2* (19) | this study |
| 50 | *hel2Δ*, *new1Δ*::KanMX, *ski2Δ*::*LEU2*, pGTSTR | BY4741, *hel2Δ*, *new1Δ*::KanMX, *ski2Δ*::*LEU2*, pGTSTR | *hel2Δ*, *new1Δ*::KanMX, *ski2Δ*::*LEU2* (20) | this study |
| 51 | *slh1Δ*::KanMX, *ski2Δ*::*LEU2*, pGTSTR | BY4741, *slh1Δ*::KanMX, *ski2Δ*::*LEU2*, pGTSTR | *slh1Δ*::KanMX, *ski2Δ*::*LEU2* (21) | this study |
| 52 | *slh1Δ*::KanMX, *new1Δ*::HphMX, *ski2Δ*::*LEU2*, pGTSTR | BY4741, *slh1Δ*::KanMX, *new1Δ*::HphMX, *ski2Δ*::*LEU2*, pGTSTR | *slh1Δ*::KanMX, *new1Δ*::HphMX, *ski2Δ*::*LEU2* (22) | this study |
| 53 | *rqt4Δ*::KanMX, *ski2Δ*::*LEU2*, pGTSTR | BY4741, *rqt4Δ*::KanMX, *ski2Δ*::*LEU2*, pGTSTR | *rqt4Δ*::KanMX, *ski2Δ*::*LEU2* (23) | this study |
| 54 | *rqt4Δ*::KanMX, *new1Δ*::HphMX, *ski2Δ*::*LEU2*, pGTSTR | BY4741, *rqt4Δ*::KanMX, *new1Δ*::HphMX, *ski2Δ*::*LEU2*, pGTSTR | *rqt4Δ*::KanMX, *new1Δ*::HphMX, *ski2Δ*::*LEU2* (24) | this study |
| 55 | *cue3Δ*::KanMX, *ski2Δ*::*LEU2*, pGTSTR | BY4741, *cue3Δ*::KanMX, *ski2Δ*::*LEU2*, pGTSTR | *cue3Δ*::KanMX, *ski2Δ*::*LEU2* (25) | this study |
| 56 | *cue3Δ*::KanMX, *new1Δ*::HphMX, *ski2Δ*::*LEU2*, pGTSTR | BY4741, *cue3Δ*::KanMX, *new1Δ*::HphMX, *ski2Δ*::*LEU2*, pGTSTR | *cue3Δ*::KanMX, *new1Δ*::HphMX, *ski2Δ*::*LEU2* (26) | this study |
| 57 | *dom34Δ*::KanMX, *ski2Δ*::*LEU2*, pGTSTR | BY4741, *dom34Δ*::KanMX, *ski2Δ*::*LEU2*, pGTSTR | *dom34Δ*::KanMX, *ski2Δ*::*LEU2* (29) | this study |
| 58 | *new1Δ*::HphMX, *dom34Δ*::KanMX, *ski2Δ*::*LEU2*, pGTSTR | BY4741, *new1Δ*::HphMX, *dom34Δ*::KanMX, *ski2Δ*::*LEU2*, pGTSTR | *new1Δ*::HphMX, *dom34Δ*::KanMX, *ski2Δ*::*LEU2* (30) | this study |
| 59 | *cue2Δ*::KanMX, *ski2Δ*::*LEU2*, pGTSTR | BY4741, *cue2Δ*::KanMX, *ski2Δ*::*LEU2*, pGTSTR | *cue2Δ*::KanMX, *ski2Δ*::*LEU2* (27) | this study |
| 60 | *new1Δ*::KanMX, *cue2Δ*::HphMX, *ski2Δ*::*LEU2*, pGTSTR | BY4741, *new1Δ*::KanMX, *cue2Δ*::HphMX, *ski2Δ*::*LEU2*, pGTSTR | *new1Δ*::KanMX, *cue2Δ*::HphMX, *ski2Δ*::*LEU2* (28) | this study |
| 61 | BY4741, pGTRR | BY4741, BY4741, pGTRR | BY4741 (1) | this study |
| 62 | *new1Δ*::KanMX, pGTRR | BY4741, *new1Δ*::KanMX, pGTRR | *new1Δ*::KanMX (2) | this study |
| 63 | *new1Δ*::HphMX, pGTRR | BY4741, *new1Δ*::HphMX, pGTRR | *new1Δ*::HphMX (3) | this study |
| 64 | *hel2Δ*, pGTRR | BY4741, *hel2Δ*, pGTRR | *hel2Δ* (4) | this study |
| 65 | *hel2Δ*, *new1Δ*::KanMX, pGTRR | BY4741, *hel2Δ*, *new1Δ*::KanMX, pGTRR | *hel2Δ*, *new1Δ*::KanMX (5) | this study |
| 66 | *slh1Δ*::KanMX, pGTRR | BY4741, *slh1Δ*::KanMX, pGTRR | *slh1Δ*::KanMX (6) | this study |
| 67 | *slh1Δ*::KanMX, *new1Δ*::HphMX, pGTRR | BY4741, *slh1Δ*::KanMX, *new1Δ*::HphMX, pGTRR | *slh1Δ*::KanMX, *new1Δ*::HphMX (7) | this study |
| 68 | *rqt4Δ*::KanMX, pGTRR | BY4741, *rqt4Δ*::KanMX, pGTRR | *rqt4Δ*::KanMX (8) | this study |
| 69 | *rqt4Δ*::KanMX, *new1Δ*::HphMX, pGTRR | BY4741, *rqt4Δ*::KanMX, *new1Δ*::HphMX, pGTRR | *rqt4Δ*::KanMX, *new1Δ*::HphMX (9) | this study |
| 70 | *cue3Δ*::KanMX, pGTRR | BY4741, *cue3Δ*::KanMX, pGTRR | *cue3Δ*::KanMX (10) | this study |
| 71 | *cue3Δ*::KanMX, *new1Δ*::HphMX, pGTRR | BY4741, *cue3Δ*::KanMX, *new1Δ*::HphMX, pGTRR | *cue3Δ*::KanMX, *new1Δ*::HphMX (11) | this study |
| 72 | *dom34Δ*::KanMX, pGTRR | BY4741, *dom34Δ*::KanMX, pGTRR | *dom34Δ*::KanMX (14) | this study |

| | | | | |
|---|---|---|---|---|
| 73 | *dom34Δ*::KanMX, *new1Δ*::HphMX, pGTRR | BY4741, *dom34Δ*::KanMX, *new1Δ*::HphMX, pGTRR | *new1Δ*::HphMX, *dom34Δ*::KanMX (15) | this study |
| 74 | *cue2Δ*::KanMX, pGTRR | BY4741, *cue2Δ*::KanMX, pGTRR | *cue2Δ*::KanMX (12) | this study |
| 75 | *new1Δ*::KanMX, *cue2Δ*::HphMX, pGTRR | BY4741, *new1Δ*::KanMX, *cue2Δ*::HphMX, pGTRR | *new1Δ*::KanMX, *cue2Δ*::HphMX (13) | this study |
| 76 | *ski2Δ*::*LEU2*, pGTRR | BY4741, *ski2Δ*::*LEU2*, pGTRR | *ski2Δ*::*LEU2* (16) | this study |
| 77 | *new1Δ*::KanMX, *ski2Δ*::*LEU2*, pGTRR | BY4741, *new1Δ*::KanMX, *ski2Δ*::*LEU2*, pGTRR | *new1Δ*::KanMX, *ski2Δ*::*LEU2* (17) | this study |
| 78 | *new1Δ*::HphMX, *ski2Δ*::*LEU2*, pGTRR | BY4741, *new1Δ*::HphMX, *ski2Δ*::*LEU2*, pGTRR | *new1Δ*::HphMX, *ski2Δ*::*LEU2* (18) | this study |
| 79 | *hel2Δ*, *ski2Δ*::*LEU2*, pGTRR | BY4741, *hel2Δ*, *ski2Δ*::*LEU2*, pGTRR | *hel2Δ*, *ski2Δ*::*LEU2* (19) | this study |
| 80 | *hel2Δ*, *new1Δ*::KanMX, *ski2Δ*::*LEU2*, pGTRR | BY4741, *hel2Δ*, *new1Δ*::KanMX, *ski2Δ*::*LEU2*, pGTRR | *hel2Δ*, *new1Δ*::KanMX, *ski2Δ*::*LEU2* (20) | this study |
| 81 | *slh1Δ*::KanMX, *ski2Δ*::*LEU2*, pGTRR | BY4741, *slh1Δ*::KanMX, *ski2Δ*::*LEU2*, pGTRR | *slh1Δ*::KanMX, *ski2Δ*::*LEU2* (21) | this study |
| 82 | *slh1Δ*::KanMX, *new1Δ*::HphMX, *ski2Δ*::*LEU2*, pGTRR | BY4741, *slh1Δ*::KanMX, *new1Δ*::HphMX, *ski2Δ*::*LEU2*, pGTRR | *slh1Δ*::KanMX, *new1Δ*::HphMX, *ski2Δ*::*LEU2* (22) | this study |
| 83 | *rqt4Δ*::KanMX, *ski2Δ*::*LEU2*, pGTRR | BY4741, *rqt4Δ*::KanMX, *ski2Δ*::*LEU2*, pGTRR | *rqt4Δ*::KanMX, *ski2Δ*::*LEU2* (23) | this study |
| 84 | *rqt4Δ*::KanMX, *new1Δ*::HphMX, *ski2Δ*::*LEU2*, pGTRR | BY4741, *rqt4Δ*::KanMX, *new1Δ*::HphMX, *ski2Δ*::*LEU2*, pGTRR | *rqt4Δ*::KanMX, *new1Δ*::HphMX, *ski2Δ*::*LEU2* (24) | this study |
| 85 | *cue3Δ*::KanMX, *ski2Δ*::*LEU2*, pGTRR | BY4741, *cue3Δ*::KanMX, *ski2Δ*::*LEU2*, pGTRR | *cue3Δ*::KanMX, *ski2Δ*::*LEU2* (25) | this study |
| 86 | *cue3Δ*::KanMX, *new1Δ*::HphMX, *ski2Δ*::*LEU2*, pGTRR | BY4741, *cue3Δ*::KanMX, *new1Δ*::HphMX, *ski2Δ*::*LEU2*, pGTRR | *cue3Δ*::KanMX, *new1Δ*::HphMX, *ski2Δ*::*LEU2* (26) | this study |
| 87 | *dom34Δ*::KanMX, *ski2Δ*::*LEU2*, pGTRR | BY4741, *dom34Δ*::KanMX, *ski2Δ*::*LEU2*, pGTRR | *dom34Δ*::KanMX, *ski2Δ*::*LEU2* (29) | this study |
| 88 | *new1Δ*::HphMX, *dom34Δ*::KanMX, *ski2Δ*::*LEU2*, pGTRR | BY4741, *new1Δ*::HphMX, *dom34Δ*::KanMX, *ski2Δ*::*LEU2*, pGTRR | *new1Δ*::HphMX, *dom34Δ*::KanMX, *ski2Δ*::*LEU2* (30) | this study |
| 89 | *cue2Δ*::KanMX, *ski2Δ*::*LEU2*, pGTRR | BY4741, *cue2Δ*::KanMX, *ski2Δ*::*LEU2*, pGTRR | *cue2Δ*::KanMX, *ski2Δ*::*LEU2* (27) | this study |
| 90 | *new1Δ*::KanMX, *cue2Δ*::HphMX, *ski2Δ*::*LEU2*, pGTRR | BY4741, *new1Δ*::KanMX, *cue2Δ*::HphMX, *ski2Δ*::*LEU2*, pGTRR | *new1Δ*::KanMX, *Cue2Δ*::HphMX, *ski2Δ*::*LEU2* (28) | this study |
| 91 | BY4741, pEV-His3 | BY4741, BY4741, pEV-His3 | BY4741 (1) | this study |
| 92 | BY4741, pNew1-FLAG-His3 | BY4741, BY4741, pNew1-FLAG-His3 | BY4741 (1) | this study |
| 93 | *new1Δ*::KanMX, pEV-His3 | BY4741, *new1Δ*::KanMX, pEV-His3 | *new1Δ*::KanMX (2) | this study |
| 94 | *new1Δ*::KanMX, pNew1-FLAG-His3 | BY4741, *new1Δ*::KanMX, pNew1-FLAG-His3 | *new1Δ*::KanMX (2) | this study |
| 95 | *ski2Δ*::*LEU2*, pEV-His3 | BY4741, *ski2Δ*::*LEU2*, pEV-His3 | *ski2Δ*::*LEU2* (16) | this study |
| 96 | *ski2Δ*::*LEU2*, pNew1-FLAG-His3 | BY4741, *ski2Δ*::*LEU2*, pNew1-FLAG-His3 | *ski2Δ*::*LEU2* (16) | this study |
| 97 | *new1Δ*::KanMX, *ski2Δ*::*LEU2*, pEV-His3 | BY4741, *new1Δ*::KanMX, *ski2Δ*::*LEU2*, pEV-His3 | *new1Δ*::KanMX (2) | this study |
| 98 | *new1Δ*::KanMX, *ski2Δ*::*LEU2*, pNew1-FLAG-His3 | BY4741, *new1Δ*::KanMX, *ski2Δ*::*LEU2*, pNew1-FLAG-His3 | *new1Δ*::KanMX (2) | this study |
| 99 | New1-HTP | BY4741, New1-HTP | BY4741 (1) | this study |
| 100 | *hel2Δ,* New1-HTP | BY4741*, hel2Δ,* New1-HTP | *hel2Δ* (4) | this study |
| 101 | Hel2-HTP | BY4741, Hel2-HTP | BY4741 (1) | Ref. (1) |
| 102 | Hel2-HTP, *new1Δ*::KanMX | BY4741, Hel2-HTP, *new1Δ*::KanMX | Hel2-HTP (101) | this study |
| 103 | New1-FTP | BY4741, New1-FTP | BY4741 (1) | this study |

**Supplementary Table S2.** List of oligonucleotides used in this study.

| Name | Sequence | Usage |
|---|---|---|
| | **Strain construction** | |
| NEW1-KanMX-HphMX-F | 5'-GTAAATACAACGACAATCAGTGCTAATTCAACT CAGGATGCGGATCCCCGGGTTAATTAA-OH-3' | deletion of *NEW1* by exchange with *KanMX* or *HphMX* cassette |
| NEW1-KanMX-R | 5'-CGAAGTTAGCGAAGATAAAACACTAGCCAGTA GGCTTTCAGAATTCGAGCTCGTTTAAAC-OH-3' | deletion of *NEW1* by exchange with *KanMX* cassette |
| NEW1-in-F | 5'-GCGTCGATCTATAGTGC-OH-3' | testing deletion of *NEW1* |
| NEW1-in-R | 5'-GAGCGTCATATGCTTGTC-OH-3' | testing deletion or tagging of *NEW1* |
| NEW1-KanMX-HphMX-F | 5'-GTAAATACAACGACATCAGTGCTAATTCAACTCA GGATGCGTACGCTGCAGGTCGAC-OH-3' | deletion of *NEW1* by exchange with *KanMX* or *HphMX* cassette |
| NEW1-KanMX-HphMX-R | 5'-CGAAGTTAGCGAAGATAAAACACTAGCCAGTA GGCTTTCAATCGATGAATTCGAGCTCG-OH-3' | deletion of *NEW1* by exchange with *KanMX* or *HphMX* cassette |
| KanMX-in-F | 5'-TTAAGTGCGCAGAAAGTAAT-OH-3' | testing presence of *KanMX* cassette |
| HphMX-in-R | 5'-ACATGGGGATGTATGGGC-OH-3' | testing presence of *KanMX* or *HphMX* cassette |
| SLH1-KanMX-F | 5'-TGAGAAGTAGATCCGTACCATCAATAGCCGGC TCAAGATGCGGATCCCCGGGTTAATTAA-OH-3' | deletion of *SLH1* by exchange with *KanMX* cassette |
| SLH1-KanMX-R | 5'-TCTTTCACAAAATAATTGTTGTTTTAATTGTGTCT CACTAGAATTCGAGCTCGTTTAAAC-OH-3' | deletion of *SLH1* by exchange with *KanMX* cassette |
| SLH1-in-F | 5'-TGAGAAGTAGATCCGTACC-OH-3' | testing deletion of *SLH1* |
| SLH1-in-R | 5'-ATCGGATTAACATTATCACCTG-OH-3' | testing deletion of *SLH1* |
| RQT4-KanMX-F | 5'-GCTTAATTATTTAACCTTGGGTTTGCAGTAATTA ATAATGCGTACGCTGCAGGTCGAC-OH-3' | deletion of *RQT4* by exchange with *KanMX* cassette |
| RQT4-KanMX-R | 5'-ATCCTCTATCATATATAATATAAACTTTTACCATT CATCAATCGATGAATTCGAGCTCG-OH-3' | deletion of *RQT4* by exchange with *KanMX* cassette |
| RQT4-in-F | 5'-GGCAAAGTGAACGTAGTC-OH-3' | testing deletion of *RQT4* |
| RQT4-in-R | 5'-AAGCCAACTAAGGAAAACTC-OH-3' | testing deletion of *RQT4* |
| CUE3-KanMX-F | 5'-TCTTGGAAATTTTTATAGGATAGAATCACTA AGAATAATGCGGATCCCCGGGTTAATTAA-OH-3' | deletion of *CUE3* by exchange with *KanMX* cassette |
| CUE3-KanMX-R | 5'-ACATACGCTTGTCTATCTTGTATGCTGATGCA TTTTATCAGAATTCGAGCTCGTTTAAAC-OH-3' | deletion of *CUE3* by exchange with *KanMX* cassette |
| CUE3-in-F | 5'-CACTAGTTCTCATAAATGAGAGAC-OH-3' | testing deletion of *CUE3* |
| CUE3-in-R | 5'-CGCTTGTCTATCTTGTATGC-OH-3' | testing deletion of *CUE3* |
| CUE2-KanMX-HphMX-F | 5'-AATTATGACACCTCATTTATCGTGCATATAAGATC ATGCATAGCTTGCCTCGTCCCC-OH-3' | deletion of *CUE2* by exchange with *KanMX* or *HphMX* cassette |
| CUE2-KanMX-HphMX-R | 5'-GCCTAGCGTTTATAACCTTTGTAGAAAAATCAGA TAGGTTCGCATAGGCCACTAGTGG-OH-3' | deletion of *CUE2* by exchange with *KanMX* or *HphMX* cassette |
| CUE2-in-F | 5'-GACACTCACAGACAAGCCTCAGGGG-OH-3' | testing deletion of *CUE2* |
| DOM34-KanMX-F | 5'-TTTTGTTCAATTATCGCATTCCTATCATAGCAAAA ATATGCGGATCCCCGGGTTAATTAA-OH-3' | deletion of *DOM34* by exchange with *KanMX* cassette |
| DOM34-KanMX-R | 5'-TTTTATGTGTACATTACTTTTTTCTTACATAGTAA ATCTAGAATTCGAGCTCGTTTAAAC-OH-3' | deletion of *DOM34* by exchange with *KanMX* cassette |
| DOM34-in-F | 5'-GCGTGATGAAAGTGTACATA-OH-3' | testing deletion of *DOM34* |
| SKI2-LEU2-F | 5'-AACCTAACTCACAAAATTTACTGTACTAATACTA ATTTATGCGTATCACGAGGCCC-OH-3' | deletion of *SKI2* by exchange with *LEU2* cassette |
| SKI2-LEU2-R | 5'-CTTTTATAAACATGACTCACATTGAGAATAAATG AGCTCTGATAAGCTGTCAAACATGAG-OH-3' | deletion of *SKI2* by exchange with *LEU2* cassette |
| SKI2-in-F | 5'-GTTAATGATATCACGACGGAC-OH-3' | testing deletion of *SKI2* |
| LEU2-in-R | 5'-GAAAAAGGTATATGCGTCAGG-OH-3' | testing presence of *LEU2* cassette |
| NEW1-HTP-F | 5'-AAGGTACACCAAAACCAGTTGATACTGACGATG AAGAAGATGAGCACCATCACCATCACC-OH-3' | tagging of New1 |
| NEW1-HTP-FTP-R | 5'-AACAAACGAAGTTAGCGAAGATAAAACACTAG CCAGTAGGCTTTACGACTCACTATAGGG-OH-3' | tagging of New1 |
| NEW1-FTP-F | 5'-AAGGTACACCAAAACCAGTTGATACTGACGAT | tagging of New1 |

| | | |
|---|---|---|
| | GAAGAAGATGAGGACTACAAAGACGATG-OH-3' | |
| HTP-in-F | 5'-TATGATTGTCTCCGGG-OH-3' | validated HTP tagging |
| | **Northern Blot** | |
| SCR1-NP | 5'-ATCCCGGCCGCCTCCATCAC-OH-3' | Northern blot probe for *SCR1* |
| PGK1-NP | 5'-TAAGATGGCCAAGAATGGTCTGGTTGGG-OH-3' | Northern blot probe for *PGK1* |
| SNR190-NP | 5'-TCGTCATGGTCGAATCGGACGAGG-OH-3' | Northern blot probe for *SNR190* |
| FLAG-NP | 5'-ACTTGTCGTCATCGTCTTTGTAGTCC-OH-3' | Northern blot probe for *FLAG* |
| GPM1-NP | 5'-AGCGTCGATTGGGGGAGGTGGAAC-OH-3' | Northern blot probe for *GPM1* |
| ADH1-NP | 5'-ACCGATCTTCCAGCCCTTAAC-OH-3' | Northern blot probe for *ADH1* |
| TDH3-NP | 5'-ACCTTCTTGGCACCAGCGTC-OH-3' | Northern blot probe for *TDH3* |
| | **Cloning** | |
| New1-Xba1-F | 5'-TCGACGGATTCTAGAATGCCTCCAAAGAAGTTTAAGG-OH-3' | Amplyfing *NEW1* CDS with Xba1 recogntion overhang |
| New1-Xba1-R | 5'-CAGGTGTCTAGAACTAGTGGTCAATCCTTGTCGT CATCGTCTTTGTAGTC-OH-3' | Amplyfing *NEW1* CDS with Xba1 recogntion overhang |
| Backbone-SangerSeq-1 | 5'-ACTCGCCATTTCAAAGAATACG-OH-3' | Sanger-Sequecning of pNew1-FLAG-His3 and pEV-His3 |
| New1-SangerSeq-1 | 5'-TCGTGAGGAAATTGCAGCCG-OH-3' | Sanger-Sequecning of pNew1-FLAG-His3 |
| New1-SangerSeq-2 | 5'-CATTCTTCAATGTCGCACTTG-OH-3' | Sanger-Sequecning of pNew1-FLAG-His3 |
| New1-SangerSeq-3 | 5'-CTCAGGCAGACTGACTTTAGAAG-OH-3' | Sanger-Sequecning of pNew1-FLAG-His3 |
| New1-SangerSeq-4 | 5'-GACGGCAAACCAATATTTGCAATG-OH-3' | Sanger-Sequecning of pNew1-FLAG-His3 |
| Mut-pBS1539-His2FLAG-F | 5'-P-GACTACAAAGACGATGACGACAAGGATTATGAT ATTCCAACTACTG-OH-3' | site-directed mutagenesis of HTP tag of pBS15139 to FTP tag, including FLAG sequence |
| Mut-pBS1539-R | 5'-P-CTCCATGGATCCTCCAG-OH-3' | site-directed mutagenesis of HTP tag of pBS15139 to FTP tag |
| pBS1539-FTP-Sanger-Seq | 5'-CGCCAAGCGCGCAATTAACCC-OH-3' | Sanger-Sequencing of mutated FTP-tag in pBS15139-FTP |
| | **CRAC** | |
| L3 | 5’-rAppTGGAATTCTCGGGTGCCAAGG-ddC-3’ | 3'-linker |
| L5Ac | 5'-invddT-ACACrGrArCrGrCrUrCrUrUrCrCrGrArUrCrU rNrNrNrGrCrGrCrArGrC-OH-3' | 5'-linker, barcode underlined |
| L5Ad | 5'-invddT-ACACrGrArCrGrCrUrCrUrUrCrCrGrArUrCrU rNrNrNrCrGrCrUrUrArGrC-OH-3' | 5'-linker, barcode underlined |
| L5Bc | 5'-invddT-ACACrGrArCrGrCrUrCrUrUrCrCrGrArUrCrU rNrNrNrCrArCrUrArGrC-OH-3' | 5'-linker, barcode underlined |
| L5Bd | 5'-invddT-ACACrGrArCrGrCrUrCrUrUrCrCrGrArUrCrU rNrNrNrUrCrUrCrUrArGrC-OH-3' | 5'-linker, barcode underlined |
| L5Ca | 5'-invddT-ACACrGrArCrGrCrUrCrUrUrCrCrGrArUrCrU rNrNrNrCrUrArGrC-OH-3' | 5'-linker, barcode underlined |
| L5Da | 5'-invddT-ACACrGrArCrGrCrUrCrUrUrCrCrGrArUrCrU rNrNrNrCrGrUrGrArUrN-OH-3' | 5'-linker, barcode underlined |
| L5Db | 5'-invddT-ACACrGrArCrGrCrUrCrUrUrCrCrGrArUrCrU rNrNrNrGrCrArCrUrArN-OH-3' | 5'-linker, barcode underlined |
| L5Dc | 5'-invddT-ACACrGrArCrGrCrUrCrUrUrCrCrGrArUrCrU rNrNrNrUrArGrUrGrCrN-OH-3' | 5'-linker, barcode underlined |
| L5De | 5'-invddT-ACACrGrArCrGrCrUrCrUrUrCrCrGrArUrCrU rNrNrNrArUrCrArCrGrN-OH-3' | 5'-linker, barcode underlined |
| L5Ea | 5'-invddT-ACACrGrArCrGrCrUrCrUrUrCrCrGrArUrCrU rNrNrNrCrArCrUrGrUrN-OH-3' | 5'-linker, barcode underlined |
| L5Eb | 5'-invddT-ACACrGrArCrGrCrUrCrUrUrCrCrGrArUrCrU rNrNrNrGrUrGrArCrArN-OH-3' | 5'-linker, barcode underlined |
| L5Ec | 5'-invddT-ACACrGrArCrGrCrUrCrUrUrCrCrGrArUrCrU rNrNrNrUrGrUrCrArCrN-OH-3' | 5'-linker, barcode underlined |
| L5Ed | 5'-invddT-ACACrGrArCrGrCrUrCrUrUrCrCrGrArUrCrU | 5'-linker, barcode underlined |

| | | |
|---|---|---|
| | rNrNrNrArCrArGrUrGrN-OH-3' | |
| PRT_D01a_RT | 5'-CAAGCAGAAGACGGCATACGAGATCCACGCTT CATTCCTGGCCTTGGCACCCGAGAATTCCA-OH-3' | RT, indexing in same step, index underlined |
| PRT_D02a _RT | 5'-CAAGCAGAAGACGGCATACGAGATGATGCTAC CATTCCTGGCCTTGGCACCCGAGAATTCCA-OH-3' | RT, indexing in same step, index underlined |
| PRT | 5’-CAGACGTGTGCTCTTCCGATCT-OH-3’ | RT without indexing |
| PPCR_r _1 | 5'-CAAGCAGAAGACGGCATACGA-OH-3' | library PCR, reverse primer for indexed cDNA (30°C samples) |
| PPCR_r_2 | 5'-CAAGCAGAAGACGGCATACGAGATCGGTCTCG GCATTCCTGGCCTTGGCACCCGAGAATTCC-OH-3' | library PCR, reverse primer for non-indexed cDNA (20°C samples) |
| PPCR_f | 5'-AATGATACGGCGACCACCGAGATCTACACTC TTTCCCTACACGACGCTCTTCCGATCT-OH-3‘ | library PCR |
| Pi_index_read | 5'-GGAATTCTCGGGTGCCAAGGCCAGGAATG-OH-3' | custom index read primer (30°C samples) |
| | **Long-read sequencing** | |
| | Native adapter(NA) (ONT, SQK-NBD114.24) | |

**Supplementary Table S3.** List of plasmids used in this study.

| Plasmid | Reference |
|---|---|
| pKK148 | (3) |
| pBS1539-FTP | this study |
| pBS1539-HTP | (4) |
| pEV-His3 | This study |
| pFA6-kanMX | (5) |
| pGTRR | (6) |
| pGTSTR | (6) |
| pNew1-FLAG-His3 | This study |
| pyM18 | (7) |
| pyM20 | (7) |
| YEp181-CUP1-His-Ubi | (8) |

**Supplementary Table S4.** Materials used in this study

| Product | Company | Product number | Comment |
|---|---|---|---|
| [gamma-P32] ATP | Hartmann Analystic | SRP-501 | Northern |
| 2-Mercaptoethanol | Roth | 4227.3 | |
| 2-Mercaptoethanol | Roth | 4227.3 | |
| 2-Propanol ≥99,7% | VWR | 20842.33 | |
| Accessories Beads Zirconia/glass beads, 0,5 mm | Roth | N034.1 | |
| Acetic acid glacial ≥99.7% | VWR | 20104.334 | |
| Agar | Formedium | AGA03 | |
| Albumin, Acetylated from bovine serum | Sigma-Aldrich | B8894 | |
| Ammonium peroxydisulphate | Roth | 9592.3 | |
| Ampicillin Sodium | Formedium | AMP05 | 100 µg/mL |
| AMPure XP beads-based | AGENCOURT® | A63881 | |
| anti-alpha Tubulin antibody | Abcam | ab184970 | 1:10,000 |
| anti-Mouse (GAM)-HRP conjugate antibody | Bio-Rad Laboratories | 1705047 | 1:20,000 |
| anti-PGK1 antibody | Thermo Scientific™ | 22C5D8 | 1:10,000 |
| ATP Solution (100 mM) | Thermo Scientific™ | R0441 | |
| Blunt/TA Ligase Master Mix | New England Biolabs | M0367 | |
| Bromophenol blue sodium salt | Roth | A512.2 | |
| Complete supplement mixture | Formedium | DCS0019 | |
| Complete supplement mixture drop out -URA | Formedium | DCS0169 | |
| cOmplete™, Mini, EDTA-free Protease Inhibitor Cocktail | Roche | 4693159001 | |
| cOmplete™, Mini, EDTA-free Protease Inhibitor Cocktail | Roche | 04693159001 | |
| D(+)-Glucose | Roth | HN06.3 | |
| Deoxycholic acid, sodium salt | Sigma-Aldrich | D6750-10G | |
| di-Sodium hydrogen phosphate | Roth | P030.1 | |
| DreamTaq Green DNA Polymerase | Thermo Scientific™ | EP0712 | |
| Dynabeads™ M-280 Tosylactivated | Invitrogen™ | 14203 | |
| E. coli Poly(A) Polymerase | New England Biolabs | M0276 | |
| ECL anti Rabbit IgG HRP | Cytiva | NA934W | 1:5,000 |
| Ethanol absolute, >= 99% | Fisher Chemical | E/0600DF/F21 | |
| Ethidium bromide solution 1 % | Roth | 2218.2 | |
| Ethylendiamine tetraacetic acid disodium salt dihydrate | Roth | 8043.2 | |
| Exonuclease I | Thermo Scientific™ | EN0582 | |
| FastAP Thermosensitive Alkaline Phosphatase | Thermo Scientific™ | EF0651 | |
| G-418 DiSulphate | Formedium | G4181 | 200 µg/mL |
| GeneJET Gel Extraction Kit | Thermo Scientific™ | K0691 | |
| GeneJET Plasmid Miniprep Kit | Thermo Scientific™ | K0503 | |
| Glycerol 99% | Grüssing | 110521000 | |
| Glycine | Roth | 79.4 | |
| Guanidine hydrochloride | Roth | 0037.1 | |
| Halo TEV protease | Promega | G660B | |
| HEPES | Formedium | HEPES01 | |

| | | | |
|---|---|---|---|
| Hybond-N | GE Healthcare | RPN303N | |
| Hybond-N+ | GE Healthcare | RPN203B | |
| Hygromycin B solution | Roth | 1287.2 | 300 µg/mL |
| IgG from rabbit serum | Sigma-Aldrich | I5006 | |
| IgG Sepharose™ 6 Fast Flow | Cytiva | 17-0969-01 | |
| Imidazole | Roth | X998.3 | |
| La Taq | Takara | RR002 | |
| L-Histidin | Roth | 1696.1 | |
| Lithium acetate | Roth | 5447.2 | |
| L-Leucin | Roth | 1699.1 | |
| L-Tryptophan | Roth | 1739.1 | |
| Maxima H Minus Double-Stranded cDNA Synthesis Kit | Thermo Scientific™ | K2561 | |
| MetaPhor Agarose | BioZym | 859181 | |
| MinElute Gel Extraction Kit | Qiagen | 28604 | |
| MOPS | Roth | 6979.2 | |
| Mouse IgG (Magnetic Bead Conjugate) | Cell Signaling | 5873 | |
| Native Barcoding Kit 24 V14 | Oxford Nanopore Technologies | SQK-NBD114.24 | |
| NEBNext® Quick Ligation Module | New England Biolabs | E6056 | |
| NEBNext® Ultra™ II End Repair/dA-Tailing Module | New England Biolabs | E7546 | |
| Ni-NTA sepharose beads | Qiagen | 30210 | |
| NorthernMax™-Gly Sample Loading Dye | Invitrogen™ | AM8551 | |
| Novex™ TBE Gels, 6% | Invitrogen™ | EC6265 | |
| NP-40 Alternative | Calbiochem | 492016 | |
| NP-40-Alternative | Sigma-Aldrich | 492016-100ML | |
| NuPAGE 4-12% Bis–Tris gels | Invitrogen™ | NP0335 | |
| PageRuler™ Plus Prestained Protein Ladder, 10 to 250 kDa | Thermo Scientific™ | 26619 | |
| PEG | Sigma-Aldrich | 88276-250G-F | |
| Phosphate Buffered Saline | Formedium | PBS100L | |
| Phusion™ High-Fidelity DNA-Polymerase | Thermo Scientific™ | F530L | |
| Pierce™ ECL Western Blotting-Substrat | Thermo Scientific™ | 32106 | |
| Ponceau S (C.I. 27195) | Roth | 5938.2 | |
| Powdered milk | Roth | T145.3 | |
| Proteinase K | Roth | 7528.1 | |
| PureCube Ni-NTA MagBeads | Cubebiotech | 31201 | |
| QIAquick Gel Extraction Kit | Qiagen | 28704 | |
| Qubit™ dsDNA Quantification Assay Kit | Invitrogen™ | Q3285 | |
| RiboRuler Low Range RNA Ladder | Thermo Scientific™ | SM1831 | |
| RNace-IT Ribonuclease Cocktail | Agilent Technologies | 400720 | |
| RNAClean XP beads | AGENCOURT® | A63987 | |
| RNase A/T1 Mix | Thermo Scientific™ | EN0551 | |
| RNasin® Ribonuclease Inhibitor | Promega | N2515 | |
| ROTI®Aqua-Phenol | Roth | A980.3 | |
| ROTI®C/I | Roth | X984.2 | |

| | | |
|---|---|---|
| ROTI®Quant universal | Roth | 120.1 |
| ROTIPHORESE®Gel 30 (37.5:1) | Roth | 3029.1 |
| Salmon Sperm DNA sodium salt | Roth | 5434.1 |
| SDS pellets | Roth | 8029.4 |
| SigmaPrepTM spin column | Sigma Aldrich | SC1000 |
| Sodium acetate | Roth | 6773.1 |
| Sodium chloride | VWR | 27808.297 |
| Sodium dihydrogen phosphate dihydrate | Roth | T879.1 |
| Sodium hydroxide | VWR | VWRC28244.295 |
| Soya peptone | Formedium | VPEP01 |
| SuperScript™ III Reverse Transcriptase | Invitrogen™ | 18080093 |
| T4 DNA Ligase | Thermo Scientific™ | EL0016 |
| T4 polynucleotide kinase | Thermo Scientific™ | EK0032 |
| T4 RNA Ligase 1 (ssRNA Ligase) | New England Biolabs | M0204 |
| T4 RNA Ligase 2, truncated KQ | New England Biolabs | M0373 |
| TEMED | Roth | 2367.1 |
| TRIS | Roth | 5429.3 |
| Trisodium citrate dihydrate >= 99% | Thermo Scientific Alfa Aesar | 036439.A3 |
| Trizma® hyfrochloride solution | Sigma-Aldrich | T2319-1L |
| TRIzol™ Reagent | Invitrogen™ | 15596018 |
| Tween® 20 | Roth | 9127.2 |
| UltraPure™ Agarose | Invitrogen™ | 16500500 |
| Uracil (Formedium, #) | Formedium | DOC0212 |
| XbaI | Thermo Scientific™ | ER0681 |
| Yeast extract | Formedium | YEA03 |
| Yeast Nitrogen Base without Amino Acids | Formedium | CYN0410 |
| Yeast synthetic drop-out medium supplements | Sigma-Aldrich | Y2001-20G |
| Zirconia/Silica Beads | BioSpec | 11079105z |

**Supplementary Table S5.** Devices used in this study.

| Device | Company |
|---|---|
| Fixed angle rotor 1189-A | Hettich |
| Flexstation | Molecular Devices |
| FUSION Pulse TS | Vilber |
| Mikro 220R centrifuge | Hettich |
| Mini Hybrid 38 | H. Saur |
| MiniSeq-System | Illumina |
| Multifuge X3R Centrifuge | Thermo Scientific™ |
| NanoDrop™ 2000 | Thermo Scientific™ |
| Orbitrap Astral mass spectrometer | Thermo Scientific™ |
| Owl™ A5 system | Thermo Scientific™ |
| Qubit™ 2 Fluorometer | Invitrogen™ |
| Qubit™ 3 Fluorometer | Invitrogen™ |
| ReproSil-Pur 120 C18-AQ | Dr. Maisch GmbH |
| Sonorex Super RK 31 | Bandelin electronic |
| Swinging bucket rotor TX-750 | Thermo Scientific™ |
| Typhoon FLA9500 | GE Healthcare |
| UV Stratalinker™ 1800 | Stratagene |
| Vanquish Neo UHPLC system | Thermo Scientific™ |
| Vari-X-Link | UVO3 |
| Wet/Tank Blotting Systems | BioRad |

**Supplementary Table S6.** Barcoded 5′-linkers used for CRAC experiments.

| Strain | Temperature | Replicate | 5'-Linker |
|---|---|---|---|
| BY4741 | 30°C | 1 | L5Ad |
| Hel2-HTP | 30°C | 1 | L5Ca |
| Hel2-HTP, *new1Δ::KanMX* | 30°C | 1 | L5Ac |
| BY4741 | 30°C | 2 | L5Dc |
| Hel2-HTP | 30°C | 2 | L5De |
| Hel2-HTP, *new1Δ::KanMX* | 30°C | 2 | L5Da |
| BY4741 | 20°C | 1 | L5De |
| Hel2-HTP | 20°C | 1 | L5Ea |
| Hel2-HTP, *new1Δ::KanMX* | 20°C | 1 | L5Eb |
| New1-HTP | 20°C | 1 | L5Ed |
| New1-HTP, *hel2Δ* | 20°C | 1 | L5Ec |
| BY4741 | 20°C | 2 | L5Bc |
| Hel2-HTP | 20°C | 2 | L5Bd |
| Hel2-HTP, *new1Δ::KanMX* | 20°C | 2 | L5Da |
| New1-HTP | 20°C | 2 | L5Db |
| New1-HTP, *hel2Δ* | 20°C | 2 | L5Dc |

**Supplementary Table S7.** Overview of Nanopore-Seq libraries generated in this work.

| Strain | Run | Replicate | Treatment | Barcode | Aligned Reads |
|---|---|---|---|---|---|
| BY4741 | 2 | 1 | /- | 10 | 1,176,568 |
| BY4741 | 2 | 2 | - | 11 | 1,462,263 |
| BY4741 | 2 | 3 | - | 12 | 1,224,305 |
| *new1Δ::KanMX* | 2 | 1 | - | 13 | 1,411,906 |
| *new1Δ::KanMX* | 2 | 2 | - | 14 | 346.781 |
| *new1Δ::KanMX* | 2 | 3 | - | 15 | 1,117,958 |
| *ski2Δ::LEU2* | 1 | 1 | +PNK, +IVPA | 19 | 481.796 |
| *ski2Δ::LEU2* | 1 | 2 | +PNK, +IVPA | 20 | 582.249 |
| *ski2Δ::LEU2* | 1 | 3 | +PNK, +IVPA | 21 | 484.108 |
| *new1Δ::KanMX, ski2Δ::LEU2* | 1 | 1 | +PNK, +IVPA | 22 | 620.611 |
| *new1Δ::KanMX, ski2Δ::LEU2* | 1 | 2 | +PNK, +IVPA | 23 | 648.142 |
| *new1Δ::KanMX, ski2Δ::LEU2* | 1 | 3 | +PNK, +IVPA | 24 | 767.655 |

**Supplementary Table S8**. Different tRNAs bound by New1 in different samples, analyzed by type. Reads mapping to different alleles of identical tRNAs were combined. Dark grey: tRNAs decoding strongly affected codons, medium grey: tRNAs decoding mildly affected codons, light grey: tRNAs decoding non-affected lysine and arginine codons.

| tRNA type | New1, rep. 1 | New1, rep. 2 | New1, *hel2Δ*, rep. 1 | New1, *hel2Δ*, rep. 2 |
|---|---|---|---|---|
| tA(AGC) | 0.78% | 0.54% | 0.68% | 0.44% |
| tA(UGC) | 6.79% | 10.37% | 8.51% | 9.36% |
| tC(GCA) | 0.86% | 0.85% | 0.48% | 0.79% |
| tD(GUC) | 2.67% | 3.44% | 6.09% | 3.81% |
| tE(CUC) | 5.91% | 4.81% | 7.18% | 4.34% |
| tE(UUC) | 18.81% | 14.17% | 21.40% | 11.28% |
| tF(GAA) | 0.04% | 0.02% | 0.11% | 0.00% |
| tG(CCC) | 1.72% | 0.77% | 0.84% | 0.80% |
| tG(GCC) | 16.16% | 11.41% | 13.65% | 8.43% |
| tG(UCC) | 3.55% | 1.23% | 2.02% | 0.92% |
| tH(GUG) | 1.92% | 1.40% | 1.64% | 1.42% |
| tI(AAU) | 1.18% | 0.95% | 0.79% | 0.63% |
| tI(UAU) | 0.12% | 0.13% | 0.13% | 0.09% |
| tK(CUU) | 3.05% | 2.26% | 3.04% | 2.57% |
| tK(UUU) | 0.35% | 0.18% | 0.23% | 0.23% |
| tL(CAA) | 3.48% | 1.20% | 1.87% | 1.17% |
| tL(GAG) | 0.03% | 0.01% | 0.03% | 0.02% |
| tL(UAA) | 0.80% | 0.49% | 0.32% | 0.37% |
| tL(UAG) | 0.61% | 0.36% | 0.56% | 0.28% |
| tM(CAU) | 0.37% | 0.30% | 0.32% | 0.21% |
| tN(GUU) | 0.58% | 0.21% | 0.76% | 0.71% |
| tP(AGG) | 0.57% | 0.02% | 0.42% | 0.08% |
| tP(UGG) | 0.31% | 0.18% | 0.15% | 0.11% |
| tQ(CUG) | 1.01% | 0.24% | 0.67% | 0.34% |
| tR(ACG) | 1.04% | 0.54% | 1.01% | 0.31% |
| tR(CCG) | 1.50% | 0.66% | 1.08% | 1.07% |
| tR(CCU) | 6.20% | 28.91% | 6.54% | 37.73% |
| tR(UCU) | 5.98% | 2.40% | 4.99% | 2.72% |
| tS(AGA) | 3.86% | 5.35% | 4.58% | 2.37% |
| tS(CGA) | 0.14% | 0.07% | 0.16% | 0.18% |
| tS(GCU) | 0.80% | 0.45% | 0.91% | 0.39% |
| tS(UGA) | 2.09% | 1.80% | 1.99% | 2.39% |
| tT(AGU) | 0.59% | 0.20% | 0.49% | 0.24% |
| tT(CGU) | 0.72% | 0.85% | 0.79% | 0.78% |
| tT(UGU) | 0.27% | 0.11% | 0.27% | 0.07% |
| tT(XXX) | 0.00% | 0.00% | 0.00% | 0.00% |
| tV(AAC) | 0.91% | 0.45% | 0.76% | 0.59% |
| tV(CAC) | 0.88% | 0.72% | 0.97% | 0.72% |
| tV(UAC) | 1.30% | 1.08% | 0.64% | 0.78% |
| tW(CCA) | 0.49% | 0.07% | 0.43% | 0.14% |
| tW(UCA) | 0.00% | 0.00% | 0.02% | 0.00% |
| tX(XXX) | 0.27% | 0.15% | 0.32% | 0.13% |
| tY(GUA) | 1.29% | 0.64% | 2.17% | 0.99% |
| total tRNA in library | 20% | 16% | 14% | 19% |
